# The ubiquitin ligase HUWE1 enhances WNT signaling by antagonizing destruction complex-mediated β-catenin degradation and through a mechanism independent of β-catenin stability

**DOI:** 10.1101/2024.02.02.578552

**Authors:** Joseph K. McKenna, Yalan Wu, Praveen Sonkusre, Raj Chari, Andres M. Lebensohn

**Affiliations:** Laboratory of Cellular and Molecular Biology, Center for Cancer Research, National Cancer Institute, National Institutes of Health, Bethesda, Maryland, United States of America; Genome Modification Core, Laboratory Animal Sciences Program, Frederick National Lab for Cancer Research, Frederick, Maryland, United States of America

## Abstract

WNT/β-catenin signaling is mediated by the transcriptional coactivator β-catenin (CTNNB1). CTNNB1 abundance is regulated by phosphorylation and proteasomal degradation promoted by a destruction complex composed of the scaffold proteins APC and AXIN1 or AXIN2, and the kinases CSNK1A1 and GSK3A or GSK3B. Loss of CSNK1A1 increases CTNNB1 abundance, resulting in hyperactive WNT signaling. Previously, we demonstrated that the HECT domain ubiquitin ligase HUWE1 is necessary for hyperactive WNT signaling in HAP1 haploid human cells lacking CSNK1A1. Here, we investigate the mechanism underlying this requirement. In the absence of CSNK1A1, GSK3A/GSK3B still phosphorylated a fraction of CTNNB1, promoting its degradation. HUWE1 loss enhanced GSK3A/GSK3B-dependent CTNNB1 phosphorylation, further reducing CTNNB1 abundance. However, the reduction in CTNNB1 caused by HUWE1 loss was disproportionately smaller than the reduction in WNT target gene transcription. To test if the reduction in WNT signaling resulted from reduced CTNNB1 abundance alone, we engineered the endogenous *CTNNB1* locus in HAP1 cells to encode a CTNNB1 variant insensitive to destruction complex-mediated phosphorylation and degradation. HUWE1 loss in these cells reduced WNT signaling with no change in CTNNB1 abundance. Genetic interaction and overexpression analyses revealed that the effects of HUWE1 on WNT signaling were not only mediated by GSK3A/GSK3B, but also by APC and AXIN1. Regulation of WNT signaling by HUWE1 required its ubiquitin ligase activity. These results suggest that in cells lacking CSNK1A1, a destruction complex containing APC, AXIN1 and GSK3A/GSK3B downregulates WNT signaling by phosphorylating and targeting CTNNB1 for degradation. HUWE1 enhances WNT signaling by antagonizing this activity. Therefore, HUWE1 enhances WNT/CTNNB1 signaling through two mechanisms, one that regulates CTNNB1 abundance and another that is independent of CTNNB1 stability. Coordinated regulation of CTNNB1 abundance and an independent signaling step by HUWE1 would be an efficient way to control WNT signaling output, enabling sensitive and robust activation of the pathway.

**Author Summary:** The WNT pathway is a conserved signaling system with diverse functions in embryonic development and adult tissue homeostasis. Dysregulation of WNT signaling drives many types of cancer. Over four decades of research have revealed a great deal about how the core components of the WNT pathway regulate signaling, but much less is known about additional regulatory layers superimposed on the core signaling module. In this study we present an example of such regulation by the ubiquitin ligase HUWE1. Phosphorylation of the transcriptional co-activator β-catenin by a protein complex called the destruction complex targets β-catenin for degradation. This is considered the main regulated step in WNT signaling. We demonstrate that HUWE1 enhances WNT signaling through two distinct mechanisms. First, HUWE1 antagonizes the phosphorylation and degradation of β-catenin by the destruction complex. Second, HUWE1 enhances WNT signaling through a mechanism independent from control of β-catenin stability. The effects of HUWE1 on WNT signaling require its ubiquitin ligase activity, suggesting there is a HUWE1 substrate awaiting discovery. Our work therefore reveals a new role for HUWE1 controlling the main regulated step in WNT signaling – β-catenin phosphorylation by the destruction complex – and most likely a downstream mechanism.

## Introduction

During embryonic development and tissue homeostasis, WNT/β-catenin signaling orchestrates cellular processes that control tissue patterning and morphogenesis, cell fate specification, and stem cell self-renewal among many other functions [1, 2]. Mutations in WNT signaling pathway components can drive tumorigenesis of many cancer types, most notably colorectal cancer [3, 4]. At the heart of the WNT/β-catenin signaling pathway, the destruction complex (DC) controls the abundance of the transcriptional coactivator β-catenin (CTNNB1) by regulating its degradation through the ubiquitin/proteasome system. The DC is comprised of a set of core components, including the scaffold proteins APC and AXIN1 or AXIN2, and the kinases casein kinase 1α (CSNK1A1) and glycogen synthase kinase 3α (GSK3A) or β (GSK3B) [5]. In the absence of signals initiated by secreted WNT ligands, CSNK1A1 phosphorylates CTNNB1 at serine (S) 45 [6, 7], priming it for further sequential phosphorylation of threonine (T) 41, S37 and S33 by GSK3A and/or GSK3B [6, 8] (we refer to residues S33, S37, T41 and S45 as the CTNNB1 phosphodegron). When phosphorylated, residues S33 and S37 create a recognition site for the ubiquitin ligase complex SCF^βTrCP^ [9, 10], which ubiquitylates CTNNB1 and targets it for proteasomal degradation [5]. Therefore, CTNNB1 abundance is kept low and WNT-dependent transcriptional programs are repressed. Binding of WNT ligands to the cell surface receptors frizzled (FZD) and LDL receptor related proteins 5 (LRP5) or 6 (LRP6) triggers the recruitment of dishevelled (DVL) and at least some DC components to FZD and LRP5/6. Formation of this receptor complex, or signalosome, downregulates the DC [11, 12] and results in accumulation of non-phosphorylated CTNNB1. CTNNB1 enters the nucleus, where it forms a complex with transcription factors of the TCF/LEF family and other coactivators to drive WNT target gene transcription [13].

This description of WNT/CTNNB1 signaling omits additional regulatory mechanisms superimposed on the core pathway that control the abundance, interactions, and subcellular localization of many components of the pathway. Such additional regulatory mechanisms tune WNT responses in diverse biological contexts, expand the functional repertoire of the pathway, and may represent potential sites of therapeutic intervention in WNT-driven cancers. Classical genetic approaches have been very successful at discovering new regulation in WNT signaling [14]. In a previous study, we sought to uncover new regulatory mechanisms in WNT signaling by performing forward genetic screens in HAP1-7TGP cells, a derivative of the haploid human cell line HAP1 harboring a fluorescent reporter of WNT signaling [15]. HAP1 cells are especially well suited for genetic screens due to the presence of a single allele of most genes in their near-haploid genome, which can be disrupted by mutagenesis to generate true genetic null cells [16]. We previously reported a comprehensive set of forward genetic screens designed to identify positive, negative and attenuating regulators of WNT/CTNNB1 signaling, as well as regulators of R-spondin (RSPO) signaling and suppressors of hyperactive WNT signaling induced by loss of distinct DC components, including APC and CSNK1A1 [15]. These screens recovered hits implicated at several levels of the pathway, including WNT and RSPO reception at the plasma membrane, cytosolic signal transduction, and transcriptional regulation. Comparative analyses of the screens enabled us to infer genetic interactions based on distinct patterns of hits identified by the different screens. The screens for suppressors of hyperactive signaling induced by loss of APC or CSNK1A1 suggested potential candidates for targeting oncogenic WNT signaling.

An unexpected outcome of the *APC* and *CSNK1A1* suppressor screens was that we observed only a partial overlap between significant hits in the two screens [15]. The phenotypic selection parameters used in both screens were the same and the cell lines used for the two screens were isogenic except for the mutations in *APC* or *CSNK1A1* we introduced by CRISPR/Cas9-mediated genome editing. Therefore, we expected that the hits identified in the two suppressor screens would be the same. After all, if APC and CSNK1A1 regulate WNT/CTNNB1 signaling through a single common function in the DC phosphorylating CTNNB1, we assumed that hyperactivating the pathway by knocking out one or the other would be functionally equivalent, and the complement of downstream regulators would be shared. While there were indeed many common hits with high significance scores in both suppressor screens, including established downstream regulators of WNT/CTNNB1 signaling such as *CTNNB1* and *CREBBP*, there were also many hits unique to the *APC* suppressor or the *CSNK1A1* suppressor screen [15]. These results suggested that the hyperactive signaling state resulting from loss of these two DC components was not equivalent. We hypothesized that the difference in potential downstream regulators in the two genetic backgrounds in which the screens were conducted – *APC* knock-out (KO) or *CSNK1A1* KO – could reflect additional roles of APC or CSNK1A1 in WNT/CTNNB1 signaling beyond their shared function regulating CTNNB1 stability.

*HUWE1*, the gene encoding the eponymous ubiquitin ligase, was the most striking example of a hit that was highly significant in the *CSNK1A1* suppressor but not the *APC* suppressor screen [15]. HUWE1 is a very large, 482 kilodalton (kDa) HECT domain ubiquitin ligase that has been implicated in many cellular processes, including transcriptional regulation, DNA replication and repair, cell cycle arrest, cell adhesion, cell migration, cell proliferation and differentiation, proteotoxic stress, ribosome biogenesis, mitochondrial maintenance, autophagy, apoptosis and WNT signaling [17–20]. *HUWE1* was the third most significant hit in the *CSNK1A1* suppressor screen, surpassed only by *CTNNB1* and *CREBBP*, which encode two of the main components of the TCF/LEF transcription complex and are therefore central players in the WNT pathway [15]. However, *HUWE1* was not a significant hit in the *APC* suppressor screen (rank number 8040 out of 11022 genes with mapped gene-trap integrations), and it was not among the most significant hits in any of the screens performed in wild-type (WT) HAP1-7TGP cells, designed to identify positive regulators of WNT3A- and RSPO1-induced signaling. These results suggested that HUWE1 might be involved in a regulatory mechanism that is most evident in the CSNK1A1^KO^ genetic background (for brevity, HAP1-7TGP cell lines in which genes were disrupted will be referred to by the name of the protein encoded by the targeted gene or genes followed by a “KO” superscript). In follow-up studies, we had confirmed that HUWE1 loss reduced WNT target gene transcription – and to a smaller extent CTNNB1 abundance – in CSNK1A1^KO^ but not in APC^KO^ cells [15]. We had also shown that microinjection of *HUWE1* mRNA into *Xenopus laevis* embryos promoted body axis duplication, a hallmark of ectopic WNT signaling [15]. These experiments established a few biological contexts in which HUWE1 acts as a positive regulator of WNT/CTNNB1 signaling, but the underlying mechanism remained unclear and the reason why HUWE1 loss selectively reduced WNT/CTNNB1 signaling in CSNK1A1^KO^ cells remained unknown.

Here we extend our genetic analyses to show that HUWE1 enhances WNT/CTNNB1 signaling through two different mechanisms. First, HUWE1 reduces phosphorylation of the CTNNB1 phosphodegron by antagonizing the activity of a DC composed of GSK3A/GSK3B, APC and AXIN1, therefore increasing CTNNB1 abundance. Second, HUWE1 enhances WNT signaling through a mechanism that is independent from the control of CTNNB1 stability.

## Results

### HUWE1 enhances WNT signaling in CSNK1A1^KO^ cells by antagonizing GSK3A/GSK3B-dependent phosphorylation of the CTNNB1 phosphodegron and increasing CTNNB1 abundance

We previously reported that HUWE1 loss in CSNK1A1^KO^ cells caused a substantial, 80-90% reduction in WNT reporter activity and endogenous WNT target gene expression that was accompanied by a smaller, 20-32% reduction in soluble CTNNB1 abundance [15]. Soluble CTNNB1 is a proxy for the signaling CTNNB1 pool because it excludes the more stable, plasma membrane-associated junctional CTNNB1 pool. We readily reproduced these results in the current study: HUWE1 loss in CSNK1A1^KO^ cells reduced WNT reporter activity by 89% and soluble CTNNB1 abundance by 36% (Figs 1A and 1B, and S1A Fig). These results raised the possibility that in CSNK1A1^KO^ cells, HUWE1 loss reduces WNT signaling solely by reducing CTNNB1 abundance, but that a non-linear relationship between changes in CTNNB1 abundance and transcriptional activity results in a disproportionately greater reduction in WNT target gene expression than CTNNB1 abundance. Alternatively, HUWE1 could regulate both CTNNB1 abundance and another process, which when disrupted together following HUWE1 loss result in a greater reduction in WNT target gene expression than in CTNNB1 abundance. To distinguish between these possibilities, we thought it was important to first determine the mechanism underlying the reduction in CTNNB1 abundance caused by HUWE1 loss.

**Fig 1.**
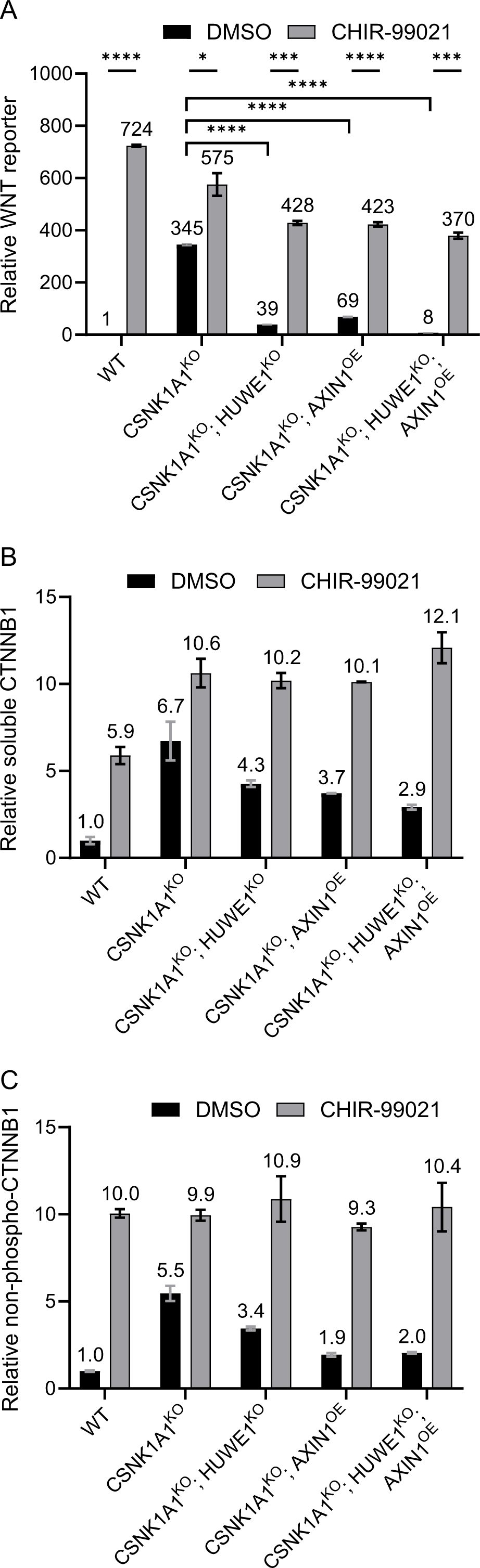
HUWE1 and AXIN1 reciprocally regulate WNT signaling by modulating GSK3A/GSK3B-dependent CTNNB1 phosphorylation and abundance. (A-C). We note that the data for WT HAP-7TGP, CSNK1A1^KO^ and CSNK1A1^KO^; HUWE1^KO^ cells is discussed in the first section of the results, while the data for CSNK1A1^KO^; AXIN1^OE^ and CSNK1A1^KO^; HUWE1^KO^; AXIN1^OE^ cells is discussed in a later section of the results subtitled “HUWE1 enhances WNT signaling by antagonizing the DC.” Cells were treated with DMSO vehicle or 10 µM of the GSK3A/GSK3B inhibitor CHIR-99021 for 48 hr as indicated. (A) WNT reporter activity (median EGFP fluorescence from 10,000 singlets was measured for triplicate wells and the average ± standard deviation (SD) of the three measurements is depicted), relative to WT HAP1-7TGP cells treated with DMSO. Significance was determined by unpaired t-test with Welch’s correction. (B) Soluble CTNNB1 abundance (CTNNB1 intensity normalized to total protein, average ± SD from duplicate immunoblots shown in S1A Fig) in membrane-free supernatants (MFS) of the indicated cell lines, relative to WT HAP1-7TGP cells treated with DMSO. (C) Non-phospho-CTNNB1 (S33/S37/T41) abundance (non-phospho-CTNNB1 intensity normalized to total protein, average ± SD from duplicate immunoblots shown in S1B Fig) in whole cell extracts (WCE) of the indicated cell lines, relative to WT HAP1-7TGP cells treated with DMSO.

The main mechanism regulating CTNNB1 abundance is phosphorylation of the CTNNB1 phosphodegron by the DC [5]. In CSNK1A1^KO^ cells we did not expect the phosphodegron to be phosphorylated by GSK3A/GSK3B at residues S33, S37 and T41 because phosphorylation of these residues generally requires the priming phosphorylation of residue S45 by CSNK1A1 [6, 7]. Nevertheless, we tested whether the reduction in CTNNB1 abundance caused by HUWE1 loss in CSNK1A1^KO^ cells was due to changes in phosphorylation of the CTNNB1 phosphodegron. CTNNB1 phosphorylated at S33, S37 and T41 can be measured directly by Western blot, but due to the rapid proteasomal degradation of this species, treatment with proteasome inhibitors is usually required to make such measurements [7]. Since any effects of HUWE1 on WNT signaling could conceivably also depend on proteasomal degradation of a HUWE1 substrate, which would be disrupted by proteasome inhibitors, we opted for a different way to evaluate phosphorylation of the CTNNB1 phosphodegron. We instead quantified CTNNB1 that is *not* phosphorylated at residues S33, S37 and T41 (we refer to this species as non-phospho-CTNNB1, but it is also known as active CTNNB1 [21]) from whole cell extracts (WCE) (Figs 1C and S1B Fig). As a control, we also measured total CTNNB1 from WCE (S1B and S1C Figs). Non-pospho-CTNNB1 abundance in the various conditions tested was different from and exhibited larger changes than total CTNNB1 abundance (Fig 1C, and S1B and S1C Figs). This indicated that non-phospho-CTNNB1 only represents a fraction of the total CTNNB1 in WCE and is likely to accurately reflect changes in phosphorylation of the CTNNB1 phosphodegron.

HUWE1 loss in CSNK1A1^KO^ cells reduced non-phospho-CTNNB1 abundance by 37%, a reduction that correlated closely with the 36% reduction in soluble CTNNB1 abundance caused by HUWE1 loss in the same cell line (Figs 1B and 1C, and S1A and S1B Figs). This correlation suggested that the reduction in CTNNB1 abundance caused by HUWE1 loss was due to increased CTNNB1 phosphorylation at S33, S37 and T41, presumably mediated by GSK3A/GSK3B. If this were the case, inhibiting GSK3A/GSK3B should reverse the reduction in both soluble and non-phospho-CTNNB1 abundance caused by HUWE1 loss. Treatment of CSNK1A1^KO^; HUWE1^KO^ cells with the GSK3A/GSK3B inhibitor CHIR-99021 indeed increased the abundance of soluble CTNNB1 by 2.4-fold and the abundance of non-phospho-CTNNB1 by 3.2-fold (Figs 1B and 1C, and S1A and S1B Figs), entirely reversing the reductions caused by HUWE1 loss. Furthermore, GSK3A/GSK3B inhibition in CSNK1A1^KO^; HUWE1^KO^ cells increased WNT reporter activity by 10.9-fold, restoring signaling to a comparable level to that in DMSO vehicle-treated CSNK1A1^KO^ cells (Fig 1A). These results indicate that even in the absence of CSNK1A1, phosphorylation of residues S33, S37 and T41 by GSK3A/GSK3B can regulate CTNNB1 abundance, and that HUWE1 loss reduces CTNNB1 abundance and WNT signaling by promoting the phosphorylation of these residues.

Since HUWE1 loss in CSNK1A1^KO^ cells increased GSK3A/GSK3B-dependent phosphorylation of the CTNNB1 phosphodegron, we wondered whether in CSNK1A1^KO^ cells containing HUWE1, residues S33, S37 and T41 in the phosphodegron might be partially phosphorylated by GSK3A/GSK3B despite the absence of CSNK1A1. CSNK1A1^KO^ cells had a relatively high abundance of soluble and non-phospho-CTNNB1, as well as high WNT reporter activity, compared to basal levels in unstimulated WT HAP1-7TGP cells (Figs 1A-C, and S1A and S1B Figs). However, GSK3A/GSK3B inhibition with CHIR-99021 in CSNK1A1^KO^ cells increased the abundance of soluble CTNNB1 by 1.6-fold and the abundance of non-phospho-CTNNB1 by 1.8-fold (Figs 1B and 1C, and S1A and S1B Figs). WNT reporter activity also increased 1.7-fold following treatment of CSNK1A1^KO^ cells with CHIR-99021 (Fig 1A). Therefore, in the absence of CSNK1A1, residual GSK3A/GSK3B-dependent phosphorylation of the CTNNB1 phosphodegron can still take place. This is presumably followed by ubiquitylation and proteasomal degradation of phosphorylated CTNNB1.

In summary, in CSNK1A1^KO^ cells, CTNNB1 is still phosphorylated by GSK3A/GSK3B at residues S33, S37 and S41 in the CTNNB1 phosphodegron, and the reduction in soluble CTNNB1 abundance caused by HUWE1 loss is due to increased GSK3A/GSK3B-dependent phosphorylation of these residues. We conclude that when present, HUWE1 antagonizes the GSK3A/GSK3B-dependent phosphorylation and ensuing degradation of CTNNB1, thereby increasing CTNNB1 abundance and promoting WNT signaling.

Our results raise two important questions. First, is control of CTNNB1 phosphorylation and abundance the only mechanism whereby HUWE1 enhances WNT signaling, or is there another mechanism distinct from the control of CTNNB1 stability? Second, is the GSK3A/GSK3B-dependent regulation of CTNNB1 abundance by HUWE1, and potentially any other mechanisms by which HUWE1 enhances WNT signaling, also mediated by other components of the DC in addition to GSK3A/GSK3B? We addressed both these questions.

### HUWE1 enhances WNT signaling through a mechanism independent of CTNNB1 stability

We next sought to determine if HUWE1 could promote WNT signaling through additional mechanisms distinct from control of CTNNB1 phosphorylation and abundance. Mutations in the CTNNB1 phosphodegron that prevent phosphorylation by CSNK1A1 and GSK3A/GSK3B render CTNNB1 insensitive to degradation by the DC [6, 7, 22]. We reasoned that introducing such mutations into the single *CTNNB1* allele of HAP1-7TGP cells would enable us to decouple control of CTNNB1 abundance from any other mechanism by which HUWE1 enhances WNT signaling.

We used CRISPR/Cas9-induced homology directed repair (HDR) to edit the codons encoding CSNK1A1 and GSK3A/GSK3B phosphorylation sites in the phosphodegron of the single endogenous *CTNNB1* locus in HAP1-7TGP cells. We introduced mutations encoding alanine (A) substitutions in the codon encoding S45, which is phosphorylated by CSNK1A1, and in the codons encoding T41 and S37, which are sequentially phosphorylated by GSK3A/GSK3B (S1 File and S2A Fig). We were unable to mutate S33, the third GSK3A/GSK3B target site.

However, recognition of CTNNB1 by SCF^βTrCP^ requires phosphorylation of both S33 and S37 [9, 10], and therefore the mutations we introduced still prevented DC-dependent CTNNB1 degradation, as we demonstrate below. We called the resulting HAP1-7TGP derivative cell line CTNNB1^ST-A^. The mutations in the *CTNNB1* locus of CTNNB1^ST-A^ cells indeed increased soluble CTNNB1 abundance 42-fold compared to unstimulated WT HAP1-7TGP cells (Fig 2A and S2B Fig), and promoted constitutive WNT signaling as judged by WNT reporter activity and endogenous WNT target gene (*AXIN2* [23], *RNF43* [23], *NKD1* [24], *TNFRSF19* [25]) expression (Figs 2B-F). Furthermore, CTNNB1 abundance, WNT reporter activity and WNT target gene expression in CTNNB1^ST-A^ cells were substantially higher than in WT HAP1-7TGP cells treated with a near-saturating dose of WNT3A conditioned media (CM) (Figs 2A-F, and S2B Fig). Stimulation of CTNNB1^ST-A^ cells with WNT3A CM did not significantly increase total CTNNB1 abundance or WNT target gene expression (S2C-E Figs). These results confirmed that the mutations we introduced into CTNNB1^ST-A^ cells rendered CTNNB1 insensitive to degradation by the DC, and therefore abolished the control of CTNNB1 stability by WNT ligands.

**Fig 2.**
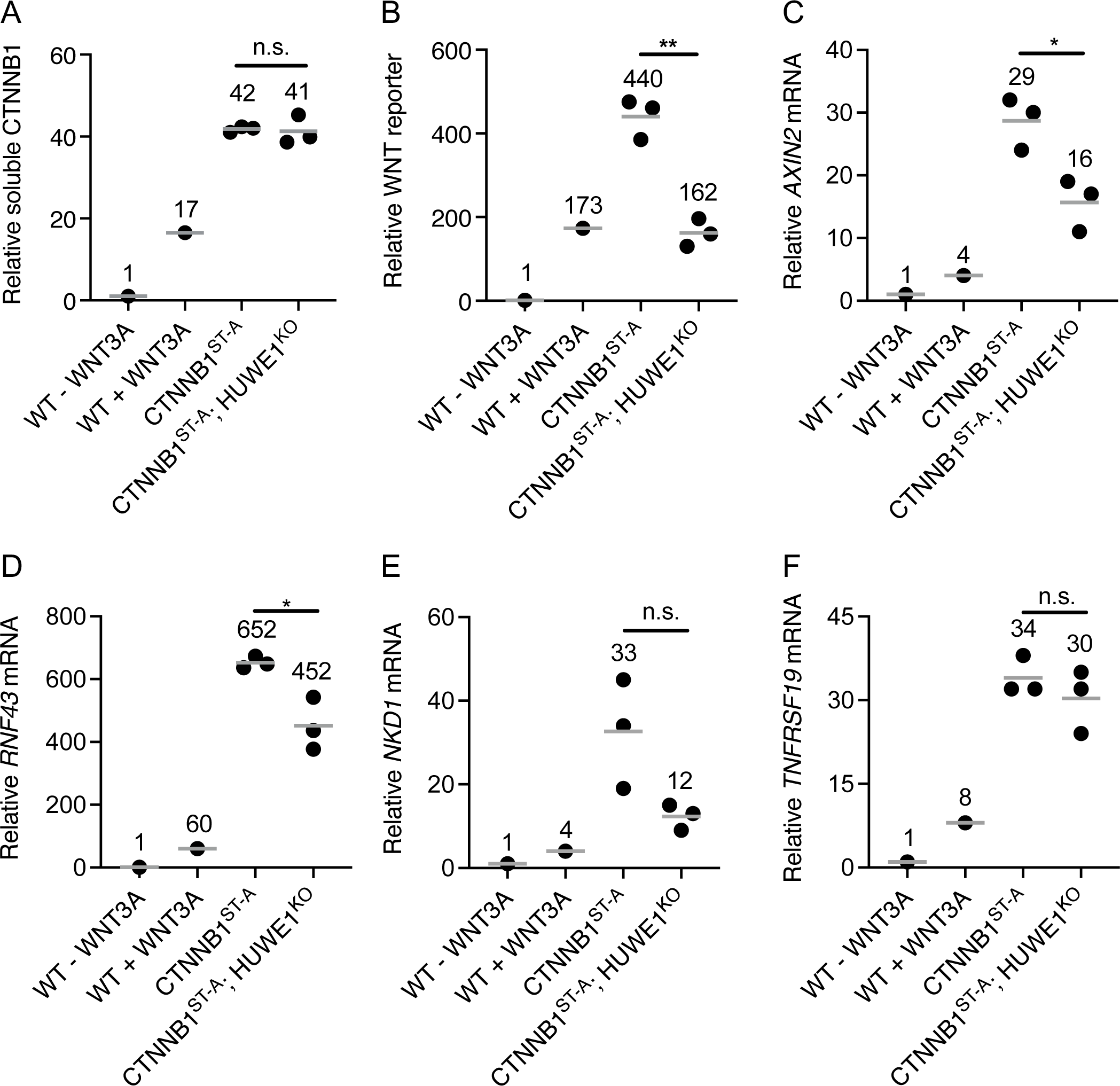
HUWE1 enhances WNT signaling through a mechanism independent of CTNNB1 stability. (A-F) Each circle represents a unique clonal cell line (determined by genotyping, S1 File). The same cell lines were used in A-F. A single value for the parental WT HAP1-7TGP cell line, and the average value from 3 independent clonal cell lines for each of the other genotypes, all relative to the untreated WT HAP1-7TGP sample, are indicated by a horizontal line and quantified above each group of circles. WT HAP1-7TGP cells were treated with 50% WNT3A CM for 24 hr where indicated. Significance was determined by unpaired t-test with Welch’s correction. (A) Relative soluble CTNNB1 abundance (CTNNB1 intensity normalized to total protein and GAPDH intensity, average from duplicate immunoblots shown in S2B Fig) in MFS of the indicated cell lines. (B) Relative WNT reporter activity (median EGFP fluorescence from 100,000 singlets). (C-F) Relative WNT target gene expression (average quantification of *AXIN2*, *RNF43*, *TNFRSF19*, or *NKD1* mRNA normalized to *HPRT1* mRNA, each measured in triplicate reactions).

We then knocked out *HUWE1* in CTNNB1^ST-A^ cells (S1 File and S2B Fig) and measured the effect on CTNNB1 abundance and WNT signaling. HUWE1 loss in multiple independent clonal cell lines (CTNNB1^ST-A^; HUWE1^KO^) did not affect soluble CTNNB1 abundance (Fig 2A and S2B Fig), but significantly reduced WNT reporter activity (Fig 2B and S2F Fig) and the expression of some WNT target genes (Figs 2C-F). These results demonstrate that HUWE1 loss reduces WNT signaling in part through a mechanism independent from the control of CTNNB1 stability. We also note that the 49% reduction in WNT reporter activity, 45% reduction in *AXIN2* expression and 31% reduction in *RNF43* expression caused by HUWE1 loss in CTNNB1^ST-A^ cells (Figs 2B-D and S2F Fig) were smaller than the 89% reduction in WNT reporter activity, 67% reduction in *AXIN2* expression and 73% reduction in *RNF43* expression caused by HUWE1 loss in CSNK1A1^KO^ cells (Figs 1A and 3C-D). This difference could be because in CSNK1A1^KO^ cells, HUWE1 loss caused a 31-36% reduction in soluble CTNNB1 abundance (Figs 1B and 3A-B, and S1A Fig) in addition to the reduction in signaling caused by the second regulatory mechanism that is independent from changes in CTNNB1 abundance, whereas no corresponding reduction in CTNNB1 abundance was observed following HUWE1 loss in CTNNB1^ST-A^ cells (Fig 2A and S2B Fig).

**Fig 3.**
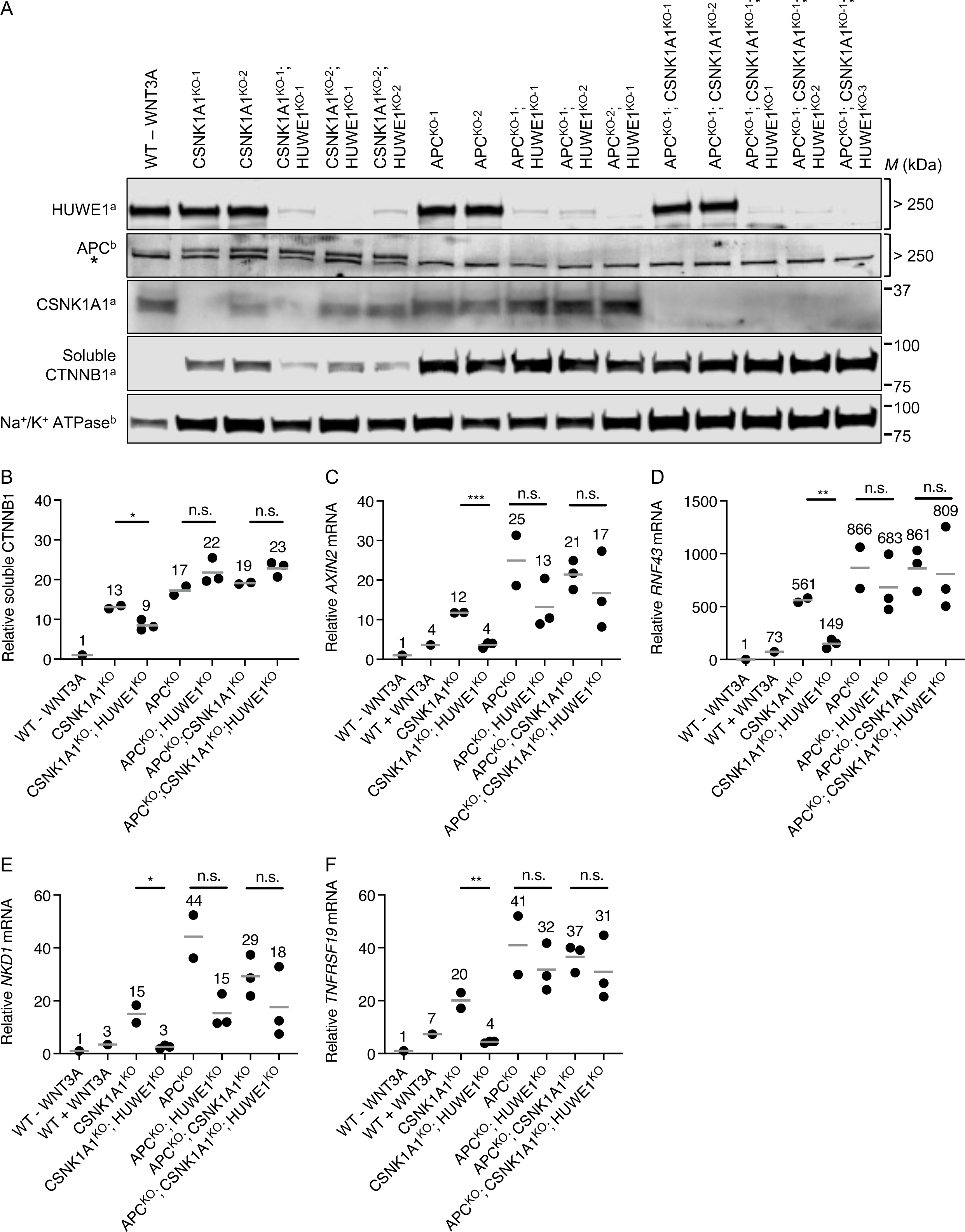
HUWE1 enhances WNT signaling through mechanisms mediated by APC. (A-F) The same cell lines were used in A-F. WT cells were treated with 50% WNT3A CM for 24 hr where indicated. (A) Immunoblot analysis of soluble proteins from MFS of the indicated clonal cell lines. We note that CSNK1A1^KO-2^ cells contain a loss-of-function mutation resulting in a 2-amino acid deletion (S1 File), and hence the protein product is still present. The “a” and “b” superscripts next to the protein names indicate which of two membranes the corresponding strips were cut from. * indicates a non-specific band observed with the mouse anti-APC antibody. (B-F) Each circle represents a unique clonal cell line (determined by genotyping, S1 File). A single value for the parental WT HAP1-7TGP cell line, and the average value from 2-3 independent clonal cell lines for each of the other genotypes, all relative to the untreated WT HAP1-7TGP sample, are indicated by a horizontal line and quantified above each group of circles. Significance was determined by unpaired t-test with Welch’s correction. (B) Relative soluble CTNNB1 abundance (CTNNB1 intensity normalized to total protein, average from duplicate immunoblots) in MFS of the indicated cell lines. (C-F) Relative WNT target gene expression (average quantification of *AXIN2*, *RNF43*, *TNFRSF19*, or *NKD1* mRNA normalized to *HPRT1* mRNA, each measured in triplicate reactions).

In summary, we distinguished two mechanisms whereby HUWE1 loss reduces WNT signaling. In CSNK1A1^KO^ cells containing WT CTNNB1, HUWE1 loss caused a moderate reduction in CTNNB1 abundance and a comparable increase in GSK3A/GSK3B-dependent phosphorylation of the CTNNB1 phosphodegron, as well as a much larger GSK3A/GSK3B-dependent reduction in WNT reporter activity (Fig 1). In CTNNB1^ST-A^ cells containing WT CSNK1A1 but a mutated CTNNB1 phosphodegron, HUWE1 loss did not alter CTNNB1 abundance but still caused a significant reduction in WNT reporter activity and WNT target gene expression (Fig 2). We conclude that HUWE1 enhances WNT signaling through two distinct mechanisms, one that increases CTNNB1 abundance and one that is independent of CTNNB1 stability.

### HUWE1 enhances WNT signaling through mechanisms mediated by APC

Having defined two mechanisms whereby HUWE1 regulates WNT signaling, a GSK3A/GSK3B-dependent mechanism that controls CTNNB1 phosphorylation and abundance, and another mechanism that is independent from the control of CTNNB1 stability, we wondered whether these mechanisms were also mediated by other DC components. HUWE1 was one of the most significant hits in a *CSNK1A1* suppressor screen but was not a significant hit in an *APC* suppressor screen [15]. Consistent with the results of these screens, HUWE1 loss substantially reduced WNT reporter activity in CSNK1A1^KO^ cells but did not affect WNT reporter activity in APC^KO^ cells [15]. Based on these results, we hypothesized that APC may be required to mediate the effects of HUWE1 on WNT signaling. If APC is required for the reduction in WNT signaling caused by HUWE1 loss in CSNK1A1^KO^ cells, then eliminating APC function in CSNK1A1^KO^; HUWE1^KO^ cells, like inhibiting GSK3A/GSK3B activity (Fig 1), should reverse said reduction. To the best of our knowledge, there are no pharmacological inhibitors that we could use to acutely inhibit APC. We were also unable to knock out *APC* in CSNK1A1^KO^; HUWE1^KO^ cells, as we found that knocking out additional genes by CRISPR/Cas9-mediated genome editing in this cell line yielded very few viable clones. Instead, we first made cell lines lacking both APC and CSNK1A1, and then tested the effects of HUWE1 loss in these cells, comparing them to cells lacking CSNK1A1 alone.

*CSNK1A1* single KO clonal HAP1-7TGP cell lines were generated and characterized previously [15] (CSNK1A1^KO-1^ and CSNK1A1^KO-2^; we note that CSNK1A1^KO-2^ is a loss-of-function allele containing a two amino acid deletion). We generated two new *APC* single KO clonal HAP1-7TGP cell lines (APC^KO-1^ and APC^KO-2^) as well as two new *APC* and *CSNK1A1* double KO clonal HAP1-7TGP cell lines (APC^KO-1^; CSNK1A1^KO-1^ and APC^KO-1^; CSNK1A1^KO-2^) using CRISPR/Cas9-mediated genome editing. We validated these cell lines by sequencing each targeted locus (S1 File), and by Western blot analysis (Fig 3A). CSNK1A1^KO^, APC^KO^ and APC^KO^; CSNK1A1^KO^ cells all exhibited elevated soluble CTNNB1 abundance several-fold higher than unstimulated WT HAP1-7TGP cells (Figs 3A and 3B). All these clonal cell lines exhibited constitutive expression of WNT target genes several-fold higher than the level of gene expression in unstimulated WT HAP1-7TGP cells and in WT HAP1-7TGP cells stimulated with a near-saturating dose of WNT3A CM (Figs 3C-F). Consistent with our results demonstrating that in CSNK1A1^KO^ cells residual phosphorylation of the CTNNB1 phosphodegron by GSK3A/GSK3B results in some CTNNB1 degradation (Fig 1), both soluble CTNNB1 abundance and WNT target gene expression were higher in APC^KO^ and APC^KO^; CSNK1A1^KO^ cells than in CSNK1A1^KO^ cells (Figs 3A-F). These results support the notion that in HAP1 cells CSNK1A1 is partially dispensable for CTNNB1 phosphorylation by GSK3A/GSK3B.

We then knocked out *HUWE1* in CSNK1A1^KO^, APC^KO^ and APC^KO^; CSNK1A1^KO^ cells to generate three CSNK1A1^KO^; HUWE1^KO^, three APC^KO^; HUWE1^KO^ and three APC^KO^; CSNK1A1^KO^; HUWE1^KO^ clonal cell lines, which we validated by sequencing the targeted *HUWE1* locus (S1 File) and by Western blot analysis (Fig 3A). HUWE1 loss in CSNK1A1^KO^ cells substantially reduced the expression of all WNT target genes tested (Figs 3C-F) and, to a lesser extent, soluble CTNNB1 abundance (Figs 3A and 3B). In contrast, HUWE1 loss in APC^KO^ cells resulted in a variable but not statistically significant reduction in WNT target gene expression (Figs 3C-F) and did not reduce soluble CTNNB1 abundance (Figs 3A and 3B), consistent with our previous finding that HUWE1 loss in APC^KO^ cells had no effect on WNT reporter activity [15]. Finally, HUWE1 loss in APC^KO^; CSNK1A1^KO^ cells yielded equivalent results to those in APC^KO^ cells, showing no significant reduction in WNT target gene expression (Figs 3C-F) or soluble CTNNB1 abundance (Figs 3A and 3B). These results indicate that, like GSK3A/GSK3B inhibition, APC loss precludes the reduction in WNT target gene expression and CTNNB1 abundance caused by HUWE1 loss in CSNK1A1^KO^ cells. We conclude that APC mediates the effects of HUWE1 on WNT signaling.

### HUWE1 enhances WNT signaling through mechanisms mediated by a subset of DC components including APC, AXIN1 and GSK3A or GSK3B

We extended the same logic as for APC (Fig 3) to test the role of every core component of the DC in mediating the functions of HUWE1 in WNT signaling. We first knocked out components of the DC individually or in certain combinations (Table 1) so we could then test the effects of HUWE1 loss on WNT signaling in each of these mutant genetic backgrounds. While we had already established the role of GSK3A/GSK3B and APC in mediating HUWE1 functions (Figs 1 and 3), we included them in our analysis to confirm those results and, in the case of GSK3A and GSK3B, test their roles individually. We used CRISPR/Cas9-mediated genome editing to generate HAP1-7TGP clonal cell lines lacking the desired DC components (Table 1). We confirmed that each targeted genomic locus had been successfully mutated (S1 File) and that the encoded protein had been eliminated (S4A Fig).

**Table 1.**
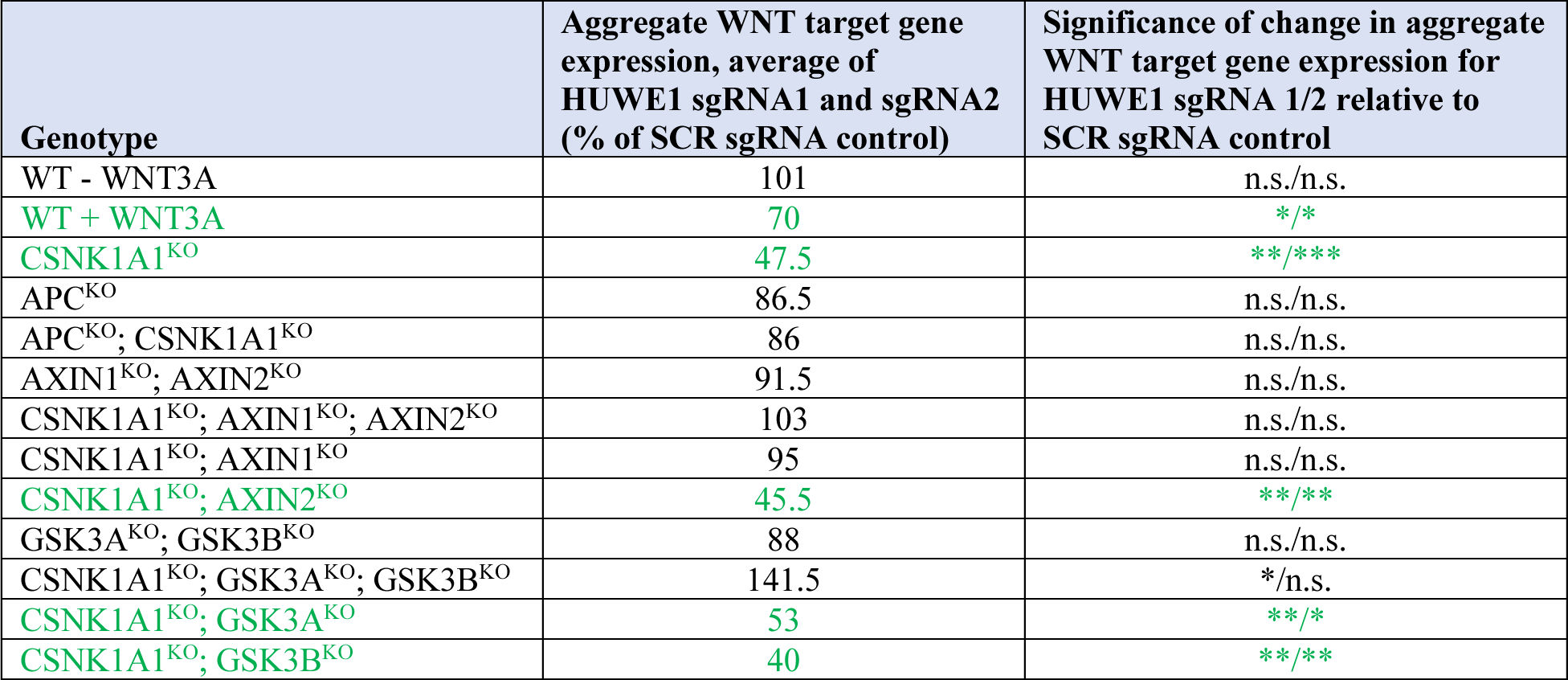
HUWE1 enhances WNT signaling through mechanisms mediated by a subset of DC components including APC, AXIN1 and GSK3A or GSK3B.

Summary of effects of CRISPRi-mediated HUWE1 KD on aggregate WNT target gene expression (Fig 4A). For the genotypes treatments indicated in green, both HUWE1 sgRNAs used resulted in a significant reduction in aggregate WNT target gene expression relative to the SCR sgRNA control.

**Fig 4.**
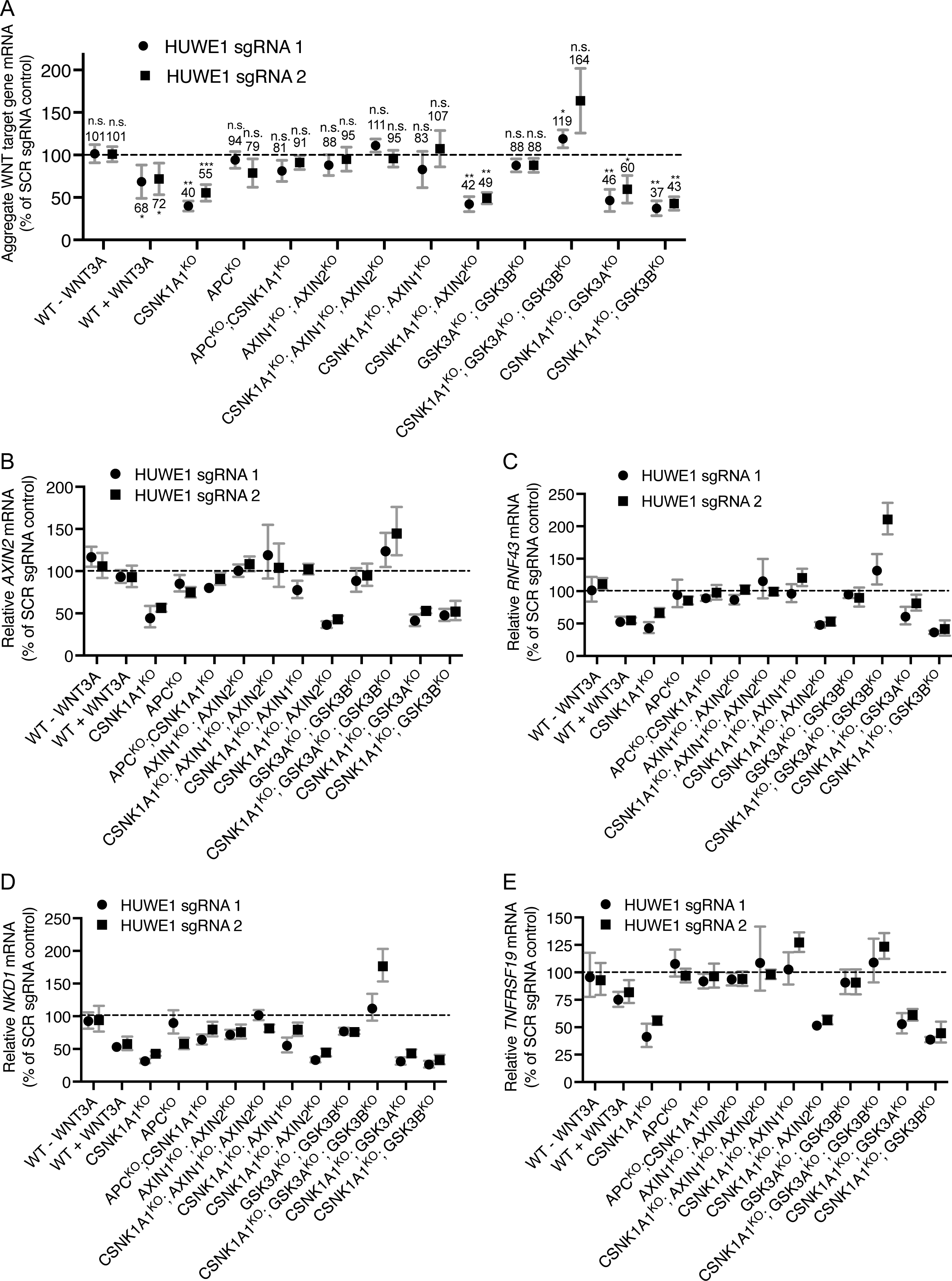
HUWE1 enhances WNT signaling through mechanisms mediated by a subset of DC components including APC, AXIN1 and GSK3A or GSK3B. (A-E) The same cell lines are used in A-E. WT HAP1-7TGP cells were treated with 50% WNT3A CM for 24 hr where indicated. (A) Aggregate WNT target gene expression (average ± SD of all four target genes, calculated from the individual average quantifications of *AXIN2*, *RNF43*, *TNFRSF19* or *NKD1* mRNA normalized to *HPRT1* mRNA, each measured in triplicate reactions) in polyclonal cell populations targeted with HUWE1 sgRNAs, reported as percentage of aggregate WNT target gene expression in polyclonal cell populations targeted with SCR sgRNA control. Significance was determined by unpaired t-test with Welch’s correction. (B-E) WNT target gene expression (average ± SD *AXIN2*, *RNF43*, *TNFRSF19*, or *NKD1* mRNA normalized to *HPRT1* mRNA, each measured in triplicate reactions) in polyclonal cell populations targeted with HUWE1 sgRNAs, reported as percentage of WNT target gene expression in polyclonal cell populations targeted with SCR sgRNA control.

To test the effects of HUWE1 loss in each of these mutant cell lines, we adopted a different experimental strategy. We had previously quantified the effect of HUWE1 loss on WNT signaling through an experimental scheme that we refer to as clonal analysis. In this scheme, we used CRISPR/Cas9-mediated genome editing to target *HUWE1*. We isolated multiple independent clonal cell lines in which *HUWE1* had been knocked out, and multiple clonal cell lines that remained WT at the targeted locus to use as controls. We then compared several KO and WT clones for WNT reporter activity or other parameters of interest (Figs 2 and 3, and S2F Fig). While clonal analysis enables comparisons in true genetic null conditions, it is subject to substantial inter-clonal variability, requiring the laborious isolation of many independent clones to achieve statistical significance. Isolation of multiple clones harboring *HUWE1* mutations in each of the 12 different genetic backgrounds (Table 1) in which we wanted to test the effect of HUWE1 loss was unfeasible. Therefore, we implemented a CRISPR interference (CRISPRi)-mediated knock-down (KD) strategy [26] that enabled us to measure the outcome of knocking down HUWE1 in polyclonal cell populations rather than in multiple individual clonal cell lines. We used a lentivirus to deliver the CRISPRi machinery together with sgRNAs targeting *HUWE1* in the various cell lines we had generated lacking DC components (Table 1 and S4A Fig). Based on Western blot measurements (S5A and S5B Figs), lentiviral delivery of either of two different sgRNAs targeting *HUWE1* (HUWE1 sgRNA1 or sgRNA2) followed by antibiotic selection of transduced cells resulted in a consistent 59-95% KD of HUWE1 compared to a control, scrambled (SCR) sgRNA. We refer to polyclonal cell populations in which we knocked down HUWE1 using CRISPRi as HUWE1^KD^, in contrast to HUWE1^KO^ clonal cell lines in which we knocked out *HUWE1* using CRISPR/Cas9-mediated genome editing.

To validate the CRISPRi KD strategy, we tested whether knocking down HUWE1 in CSNK1A1^KO^, APC^KO^, and APC^KO^; CSNK1A1^KO^ cell populations produced equivalent results to those we had observed when we knocked out HUWE1 and conducted clonal analysis in these same cell lines (Fig 3). Consistent with our clonal analysis, HUWE1 KD in CSNK1A1^KO^ cells significantly reduced the expression of four WNT target genes compared to CSNK1A1^KO^ cells transduced with SCR sgRNA (Figs 4A-E and Table 1). However, this reduction was smaller than that caused by complete HUWE1 loss in CSNK1A1^KO^; HUWE1^KO^ cells (Figs 3C-F), presumably owing to some residual HUWE1 protein present in CSNK1A1^KO^; HUWE1^KD^ cells (S5A and S5B Figs). Also consistent with our clonal analysis, HUWE1 KD in APC^KO^ and in APC^KO^; CSNK1A1^KO^ cells did not cause a statistically significant reduction in WNT target gene expression (Figs 4A-E and Table 1). These results validate CRISPRi-mediated HUWE1 KD in polyclonal cell populations as a reliable alternative to the more laborious clonal analysis of multiple individual HUWE1^KO^ clonal cell lines. We also knocked down HUWE1 in WT HAP1-7TGP cells (S5A and S5B Figs), in which we had previously reported that HUWE1 KO did not cause a significant reduction in WNT reporter activity or *AXIN2* expression induced by a near-saturating dose of WNT3A [15]. In agreement with those results, HUWE1 KD did not reduce WNT3A-induced expression of *AXIN2* (Fig 4B). However, HUWE1 KD in WT HAP1-7TGP cells did reduce the expression of other WNT target genes, including *RNF43*, *NKD1* and *TNFRSF19*, but to a smaller extent than in CSNK1A1^KO^ cells (Figs 4A-E and Table 1). Together with our analysis in CTNNB1^ST-A^ cells (Fig 2), these results demonstrate that the contribution of HUWE1 to WNT signaling is not limited to cells lacking CSNK1A1.

We then asked whether other DC components mediate the function of HUWE1. As we had done for APC, we tested whether knocking out AXIN1 and AXIN2 eliminated the reduction in WNT signaling caused by HUWE1 loss. In WT HAP1-7TGP cells, AXIN1 and AXIN2 are functionally redundant in their capacity to suppress WNT signaling, presumably by regulating CTNNB1 abundance as scaffolds in the DC: eliminating either AXIN1 or AXIN2 has no effect on WNT reporter activity, whereas eliminating both promotes constitutive pathway activation [15]. We initially assumed that a possible role of AXIN1 and AXIN2 in mediating the effects of HUWE1 may also be redundant, so we knocked out both paralogs in HAP1-7TGP cells (S1 File and S4A Fig) and tested their contribution following HUWE1 KD (S5A and S5B Figs). HUWE1 KD in AXIN1^KO^; AXIN2^KO^ cells did not reduce WNT target gene expression (Figs 4A-E and Table 1), suggesting that AXIN1, AXIN2 or both mediate the effects of HUWE1 on WNT signaling, similarly to what we had observed for APC (Fig 3). As we had done for GSK3A/GSK3B (Fig 1) and APC (Fig 3), we also tested whether the combined loss of AXIN1 and AXIN2 eliminated the reduction in WNT signaling caused by HUWE1 KD in CSNK1A1^KO^ cells. Indeed, knocking down HUWE1 in CSNK1A1^KO^; AXIN1^KO^; AXIN2^KO^ cells did not reduce WNT target gene expression as it did in CSNK1A1^KO^ cells (Figs 4A-E and Table 1). These results confirmed that AXIN1, AXIN2 or both mediate the effects of HUWE1 on WNT signaling in CSNK1A1^KO^ cells. While AXIN1 and AXIN2 are redundant in their capacity to suppress WNT signaling in WT HAP1-7TGP cells [15], it was conceivable that they may not be redundant in mediating the function of HUWE1 in CSNK1A1^KO^ cells. To test for individual contributions of AXIN1 or AXIN2, we knocked each of them out individually in CSNK1A1^KO^ cells (S1 File and S4A Fig) and then knocked down HUWE1 (S5A and S5B Figs). AXIN1 loss in CSNK1A1^KO^ cells eliminated the reduction in WNT signaling caused by HUWE1 KD, but AXIN2 loss did not (Figs 4A-E and Table 1). These results suggest that, in contrast to its redundant function with AXIN2 in suppressing WNT signaling in WT HAP1-7TGP cells [15], AXIN1 plays a unique role in mediating HUWE1-dependent effects on WNT signaling that is not redundant with AXIN2.

Given these results, we wondered whether GSK3A and GSK3B are redundant in mediating the functions of HUWE1 in WNT signaling. To answer this question, we did an equivalent series of experiments as the one we did to determine the individual roles of AXIN1 and AXIN2 in mediating HUWE1 function. Like AXIN1 and AXIN2, GSK3A and GSK3B are functionally redundant in their capacity to suppress WNT signaling in WT HAP1-7TGP cells: eliminating either GSK3A or GSK3B has no effect on WNT reporter activity, whereas eliminating both promotes constitutive pathway activation (S1 File, and S4B and S4C Figs). HUWE1 KD in GSK3A^KO^; GSK3B^KO^ cells (S5A and S5B Figs) did not reduce WNT target gene expression (Figs 4A-E and Table 1), suggesting that GSK3A, GSK3B or both mediate the effects of HUWE1 on WNT signaling. Next, we tested whether the combined loss of GSK3A and GSK3B eliminated the reduction in WNT signaling caused by HUWE1 KD in CSNK1A1^KO^ cells. Knocking down HUWE1 in CSNK1A1^KO^; GSK3A^KO^; GSK3B^KO^ cells (S5A and S5B Figs) did not reduce – and in fact increased – WNT target gene expression (Figs 4A-E and Table 1). These results confirmed that GSK3A, GSK3B or both mediate the effects of HUWE1 on WNT signaling in CSNK1A1^KO^ cells. However, unlike their combined loss, loss of GSK3A or GSK3B individually in CSNK1A1^KO^ cells did not eliminate the reduction in WNT signaling caused by HUWE1 KD (Fig 4A-E and Table 1). We conclude that the role of GSK3A and GSK3B in mediating HUWE1-dependent effects on WNT signaling is redundant, similarly to their role suppressing WNT signaling in WT HAP1-7TGP cells (S4C Fig). Therefore, only the combined loss of GSK3A and GSK3B eliminates the reduction in WNT signaling caused by HUWE1 KD in CSNK1A1^KO^ cells (Fig 4A-E and Table 1).

We considered the possibility that the distinct outcomes of knocking down HUWE1 in the various genetic backgrounds we tested (Table 1) could be due to differences in the steady state abundance of HUWE1 caused by loss of some DC complex components but not others, rather than due to other effects of distinct DC components in mediating HUWE1 function. Standard Western blot analysis did not reveal obvious differences in steady state HUWE1 abundance among the various genetic backgrounds in which we knocked down HUWE1 (S4A Fig). We corroborated this result by quantitative dot blot analysis (see Materials and methods) and did not detect significant differences in HUWE1 abundance among the different genetic backgrounds (S4D Fig).

In conclusion, a subset of DC components, including APC, AXIN1 and GSK3A or GSK3B, but not CSNK1A1 or AXIN2, mediates the function of HUWE1 in WNT signaling. Since HUWE1 enhances WNT signaling by increasing CTNNB1 abundance (Fig 1) and through another mechanism independent from the control of CTNNB1 stability (Fig 2), a DC composed of APC, AXIN1 and GSK3A/GSK3B must mediate the effects of HUWE1 on one or both mechanisms.

### HUWE1 enhances WNT signaling by antagonizing the DC

The results presented so far are consistent with the following hypothesis: 1. In CSNK1A1^KO^ cells, APC, AXIN1 and GSK3A/GSK3B are part of a DC that can partially suppress WNT signaling by phosphorylating the CTNNB1 phosphodegron and targeting CTNNB1 for proteasomal degradation; 2. HUWE1 enhances WNT signaling by antagonizing CTNNB1 phosphorylation and degradation mediated by this DC, and through another mechanism independent of CTNNB1 stability. Whether the second mechanism is also mediated by the DC remains unclear. Since all the data presented above were from loss-of-function genetic experiments, we tested this hypothesis further through overexpression experiments. Based on this hypothesis, we predicted that overexpressing the DC scaffold AXIN1 in CSNK1A1^KO^ cells should increase DC activity and therefore have similar effects as knocking out HUWE1: it should reduce WNT signaling by promoting GSK3A/GSK3B-dependent phosphorylation and degradation of CTNNB1, and possibly by promoting the second mechanism independent of CTNNB1 stability. Furthermore, since AXIN1 loss in CSNK1A1^KO^ cells eliminated the reduction in WNT signaling caused by HUWE1 loss (Figs 4A-E and Table 1), we reasoned that overexpressing AXIN1 in CSNK1A1^KO^; HUWE1^KO^ cells should have the opposite effect and synergize with HUWE1 loss to reduce WNT signaling. To test these predictions, we stably overexpressed human AXIN1 in CSNK1A1^KO^ and in CSNK1A1^KO^; HUWE1^KO^ cells through lentiviral delivery of *AXIN1* cDNA followed by antibiotic selection. We obtained polyclonal cell populations (CSNK1A1^KO^; AXIN1^OE^ and CSNK1A1^KO^; HUWE1^KO^; AXIN1^OE^, respectively) in which AXIN1 abundance was at least 2-fold higher than that in the respective parental cell lines (S1D Fig).

AXIN1 overexpression in CSNK1A1^KO^ cells indeed reduced WNT reporter activity by 80%, which was comparable to the 89% reduction caused by HUWE1 loss in CSNK1A1^KO^ cells (Fig 1A). AXIN1 overexpression combined with HUWE1 loss in CSNK1A1^KO^ cells reduced WNT reporter activity by 98%, nearly down to the basal level of unstimulated WT HAP1-7TGP cells (Fig 1A). Therefore, AXIN1 overexpression in CSNK1A1^KO^ cells phenocopied HUWE1 loss, and AXIN1 overexpression in CSNK1A1^KO^; HUWE1^KO^ cells synergized with HUWE1 loss to reduce WNT signaling. We conclude that HUWE1 and AXIN1 exert opposing effects on WNT signaling.

To test whether the reduction in WNT signaling caused by AXIN1 overexpression and by HUWE1 loss was due to the same underlying mechanisms, we measured the abundance of soluble and non-phospho-CTNNB1 in CSNK1A1^KO^; AXIN1^OE^ and CSNK1A1^KO^; HUWE1^KO^; AXIN1^OE^ cells, as we had done in WT HAP1-7TGP, CSNK1A1^KO^ and CSNK1A1^KO^; HUWE1^KO^ cells (Figs 1B and 1C, and S1A and S1B Figs). AXIN1 overexpression in CSNK1A1^KO^ cells caused a 45% reduction in soluble CTNNB1 abundance and a 64% reduction in non-phospho-CTNNB1 abundance (Figs 1B and 1C, and S1A and S1B Figs). These reductions were comparable to and greater than the respective 36% and 37% reductions caused by HUWE1 loss in CSNK1A1^KO^ cells (Figs 1B and 1C, and S1A and S1B Figs). AXIN1 overexpression combined with HUWE1 loss in CSNK1A1^KO^ cells reduced soluble CTNNB1 abundance by 57% and non-phospho-CTNNB1 by 62% (Figs 1B and 1C, and S1A and S1B Figs). These results indicate that HUWE1 and AXIN1 have opposing functions regulating a common mechanism: AXIN1 promotes CTNNB1 phosphodegron phosphorylation and the resulting reduction in CTNNB1 abundance, while HUWE1 antagonizes both.

If HUWE1 and AXIN1 exert opposing effects on WNT signaling by regulating the same GSK3A/GSK3B-dependent processes – CTNNB1 phosphorylation and abundance, and potentially another mechanism independent of CTNNB1 stability – then GSK3A/GSK3B inhibition should reverse the effects of AXIN1 overexpression in CSNK1A1^KO^ cells, as it reverses the effects of HUWE1 loss (Figs 1A-C, and S1A and S1B Figs). Therefore, we tested whether the changes in WNT reporter activity, soluble CTNNB1 abundance and CTNNB1 phosphodegron phosphorylation caused by AXIN1 overexpression alone or combined with HUWE1 loss were dependent on GSK3A/GSK3B activity. Treatment of CSNK1A1^KO^; AXIN1^OE^ cells with the GSK3A/GSK3B inhibitor CHIR-99021 reversed the effects of AXIN1 overexpression, increasing WNT reporter activity as well as soluble and non-phospho-CTNNB1 abundance to levels higher than those measured in DMSO vehicle treated-CSNK1A1^KO^ cells, and comparable to those measured in CHIR-99021-treated CSNK1A1^KO^ cells (Figs 1A-C, and S1A and S1B Figs). GSK3A/GSK3B inhibition with CHIR-99021 also reversed the synergistic reduction in WNT reporter activity, as well as the reduction in soluble and non-phospho-CTNNB1 abundance, caused by combined AXIN1 overexpression and HUWE1 loss in CSNK1A1^KO^ cells (Figs 1A-C, and S1A and S1B Figs). These results demonstrate that in CSNK1A1^KO^ cells, HUWE1 enhances and AXIN1 inhibits WNT signaling by opposing mechanisms mediated by GSK3A/GSK3B. Altogether, our results support the hypothesis that AXIN1, acting as a scaffold in the DC, promotes GSK3A/GSK3B-dependent CTNNB1 phosphorylation and degradation, even in the absence of CSNK1A1. HUWE1 enhances WNT signaling by antagonizing this DC activity.

### Regulation of WNT signaling by HUWE1 requires its ubiquitin ligase activity

HUWE1 is a very large 482 kDa ubiquitin ligase with many protein-protein interaction domains in addition to its catalytic HECT domain [27]. Therefore, it was important to determine whether the ubiquitin ligase activity of HUWE1 was required for its functions enhancing WNT signaling. HECT domain ubiquitin ligases form a covalent intermediate between a catalytic cysteine (C) residue in the HECT domain and ubiquitin before ubiquitin is transferred to the substrate [28]. We used CRISPR-mediated base editing [29] to engineer the endogenous *HUWE1* locus of CSNK1A1^KO^ cells, introducing a single point mutation that replaced the catalytic C4341 residue with arginine (R). We isolated three independent clonal cell lines (CSNK1A1^KO^; HUWE1^C4341R-1^, CSNK1A1^KO^; HUWE1^C4341R-2^ and CSNK1A1^KO^; HUWE1^C4341R-3^) in which we confirmed by sequencing that the intended point mutation had been introduced (S1 File). We compared the effects of eliminating the catalytic activity of HUWE1 to those of knocking out HUWE1 on WNT signaling. All three CSNK1A1^KO^; HUWE1^C4341R^ clonal cell lines exhibited a substantial 89-94% reduction in WNT reporter activity and a 79-88% reduction in the expression of three WNT target genes, equivalent to what we observed in CSNK1A1^KO^; HUWE1^KO^ cells (Figs 5A-D). The C4341R point mutation did not affect HUWE1 protein stability as determined by Western blot analysis of the three CSNK1A1^KO^; HUWE1^C4341R^ cell lines (Fig 5E). In contrast, no HUWE1 protein was detected in CSNK1A1^KO^; HUWE1^KO^ cells (Fig 5E). Therefore, the reduction in WNT signaling measured in CSNK1A1^KO^; HUWE1^C4341R^ cells was not due to loss of HUWE1 protein, but rather due to the elimination of its catalytic activity. We conclude that the ubiquitin ligase activity of HUWE1 is required for its functions enhancing WNT signaling.

**Fig 5.**
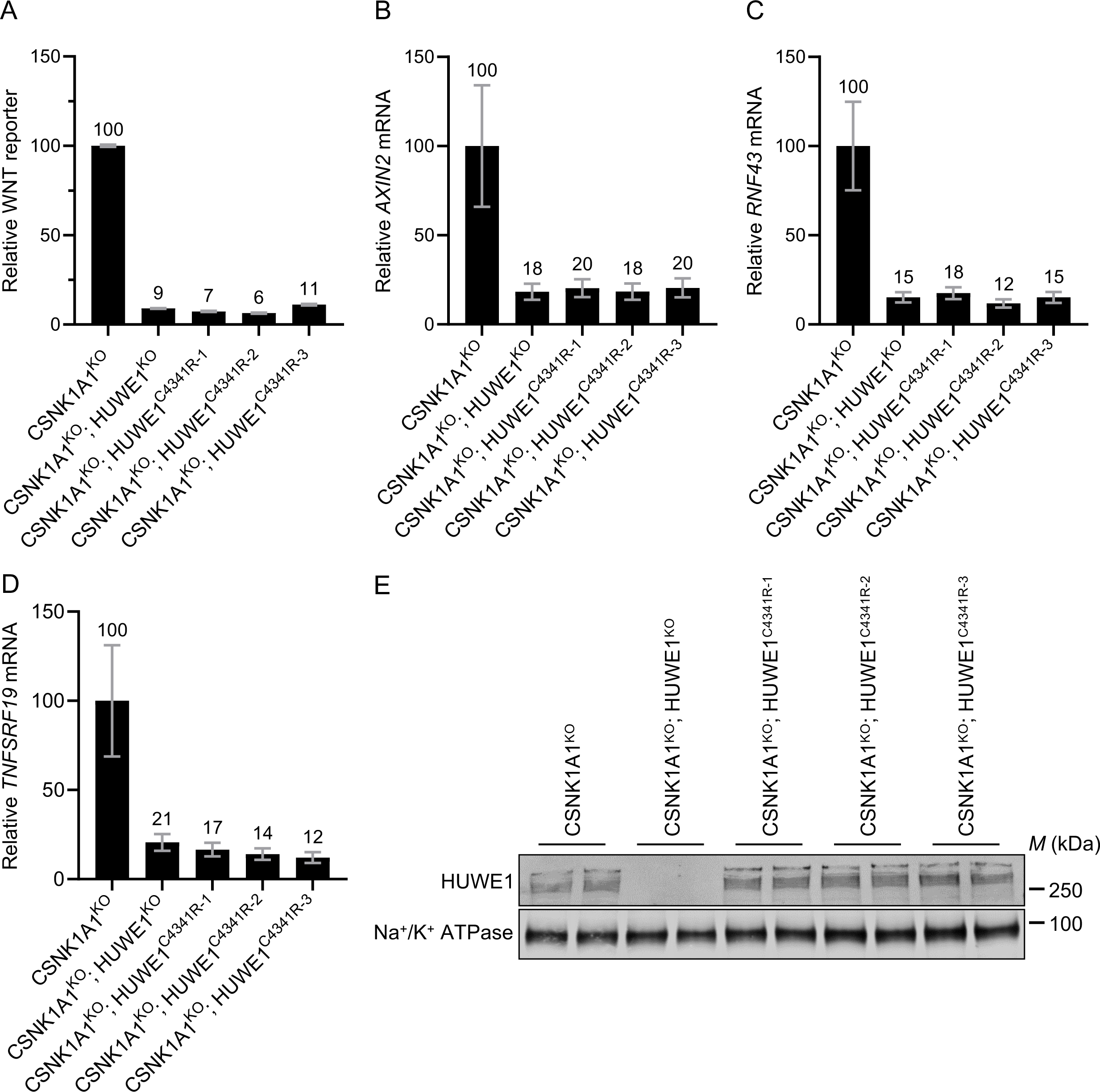
Regulation of WNT signaling by HUWE1 requires its ubiquitin ligase activity. (A) WNT reporter activity (median ± standard error of the median (SEM) EGFP fluorescence from 37,000-50,000 cells) of one CSNK1A1^KO^; HUWE1^KO^ clonal cell line and three catalytic mutant CSNK1A1^KO^; HUWE1^C4341R^ clonal cell lines, relative to CSNK1A1^KO^ cells. (B-D) WNT target gene expression (average ± SD *AXIN2*, *RNF43* or *TNFSRF19* mRNA normalized to *HPRT1* mRNA, each measured in triplicate reactions) of the indicated clonal cell lines, relative to CSNK1A1^KO^ cells. (E) Immunoblot analysis of total HUWE1 protein in WCE of the same cell lines used in A-D.

## Discussion

In this study we probed the mechanisms underlying the requirement for the HECT domain ubiquitin ligase HUWE1 to sustain hyperactive WNT/CTNNB1 signaling [15]. We demonstrate that HUWE1 enhances WNT/CTNNB1 signaling through two distinct mechanisms: by antagonizing DC-mediated CTNNB1 phosphorylation and degradation, and through another mechanism independent of CTNNB1 stability. These results are significant for two main reasons. First, they reveal a new mechanism that controls CTNNB1 stability, the main regulated step in WNT/CTNNB1 signaling. Second, by controlling another downstream step in the pathway, HUWE1 adds a new layer of regulation superimposed on the core WNT/CTNNB1 signaling module. Importantly, the coordinated regulation of CTNNB1 abundance and an independent signaling step in the pathway by HUWE1 would be an efficient way to control multiple processes that determine WNT signaling output. This may enable sensitive and robust activation of the pathway.

In CSNK1A1^KO^ cells, GSK3A/GSK3B still phosphorylate a fraction of CTNNB1 at residues S33, S37 and T41 in the phosphodegron, which reduces CTNNB1 abundance (Fig 1). HUWE1 enhances signaling by counteracting DC-dependent phosphorylation of these residues, since HUWE1 loss in CSNK1A1^KO^ cells increases phosphorylation and reduces both CTNNB1 abundance and WNT signaling activity (Fig 1). However, the reduction in CTNNB1 abundance caused by HUWE1 loss appears insufficient to account for the larger reduction in WNT target gene expression (Fig 1), suggesting that HUWE1 also enhances WNT/CTNNB1 signaling through another mechanism. In CTNNB1^ST-A^ cells containing mutations in the CTNNB1 phosphodegron that render CTNNB1 abundance insensitive to regulation by WNT ligands and the DC, HUWE1 enhances WNT target gene expression through a mechanism distinct from the control of CTNNB1 stability (Fig 2). Furthermore, regulation of WNT/CTNNB1 signaling by HUWE1 is mediated by a subset of DC components, including APC, AXIN1 and GSK3A or GSK3B, but excluding CSNK1A1 and AXIN2 (Figs 1, 3 and 4). HUWE1 promotes WNT signaling by antagonizing the activity of this DC (Fig 1). The ubiquitin ligase activity of HUWE1 is required to enhance WNT signaling (Fig 5), suggesting that a substrate of HUWE1 mediates its function.

One of the mechanisms whereby HUWE1 enhances WNT/CTNNB1 signaling is by antagonizing phosphorylation of the CTNNB1 phosphodegron by the DC complex, thereby increasing CTNNB1 abundance, but surprisingly this happens in the absence of CSNK1A1. These results demonstrate that in HAP1 cells, CSNK1A1 is not absolutely required for GSK3A/GSK3B-dependent phosphorylation of residues S33, S37 and T41 in the CTNNB1 phosphodegron, either because GSK3A/GSK3B can phosphorylate these residues without the priming phosphorylation of S45 by CSNK1A1, or because other kinases phosphorylate S45 in the absence of CSNK1A1. While priming of S45 by CSNK1A1 is generally considered a requirement for phosphorylation of S33, S37 and T41 by GSK3A/GSK3B [6, 7], some reports suggest it is not [30, 31].

We also show there is another mechanism whereby HUWE1 enhances WNT/CTNNB1 signaling that is independent of CTNNB1 stability. HUWE1 could potentially regulate CTNNB1 subcellular localization or its interactions with the TCF/LEF transcription complex, or it could regulate other downstream steps in the pathway. Elucidating this second mechanism and whether it is also mediated by a subset of DC complex components, like HUWE1-dependent regulation of CTNNB1 abundance, will be crucial to understand the full scope of how HUWE1 regulates WNT signaling.

Intriguingly, only a subset of DC components, including APC, AXIN1 and GSK3A or GSK3B, but not CSNK1A1 or AXIN2, mediate the function of HUWE1 in WNT signaling (Figs 1, 3 and 4). We were surprised to find that AXIN1 was required to mediate the effects of HUWE1 but AXIN2 was not. In WT HAP1-7TGP cells, AXIN1 and AXIN2 are redundant in their capacity to suppress WNT signaling: eliminating either AXIN1 or AXIN2 has no effect on WNT reporter activity, whereas eliminating both results in constitutive pathway activation [15]. Yet, in CSNK1A1^KO^ cells, only AXIN1 loss eliminated the reduction in WNT target gene expression caused by HUWE1 KD (Fig 4 and Table 1). These results suggest that AXIN1 and AXIN2 are not redundant in their capacity to mediate the effects of HUWE1, at least in the absence of CSNK1A1. This finding is unexpected given that mouse AXIN1 and AXIN2 proteins have been reported to be functionally equivalent *in vivo* [32], and will require further investigation.

The ubiquitin ligase activity of HUWE1 is required to promote WNT signaling in CSNK1A1^KO^ cells (Fig 5). What are the relevant ubiquitylated HUWE1 substrates, and how do they regulate WNT signaling? Does HUWE1-dependent ubiquitylation target putative substrates for proteasomal degradation or does it regulate their activity? Since a subset of DC components mediates the effects of HUWE1 on WNT signaling, is the abundance or activity of a DC component regulated by HUWE1-dependent ubiquitylation or are there other ubiquitylated substrates that indirectly impinge on DC abundance or activity? Identification of the relevant HUWE1 substrates should help answer these questions.

Previous reports have implicated HUWE1 as a *negative* regulator of WNT signaling [33–36]. This is the opposite of what we find in WT HAP1-7TGP, CSNK1A1^KO^ and CTNNB1^ST-A^ cells, in which HUWE1 is a *positive* regulator of the pathway: eliminating HUWE1 or its catalytic activity in these cells substantially reduces WNT/CTNNB1 signaling (Figs 1-5). HUWE1 has been reported to polyubiquitylate DVL and prevent DVL multimerization [33], which is required to form a functional signalosome and transduce WNT signals [37, 38]. HUWE1 has also been reported to ubiquitylate CTNNB1 and promote CTNNB1 degradation [34]. The latter mechanism is in fact the opposite of what we find in CSNK1A1^KO^ cells, in which HUWE1 loss reduces CTNNB1 abundance (Fig 1). Based on both reported mechanisms, HUWE1 loss would be expected to promote rather than reduce WNT signaling. Therefore, we do not think that DVL or CTNNB1 are the relevant ubiquitylated substrates that mediate the effects of HUWE1 on WNT signaling in HAP1 cells. These disparate results could reflect differences in experimental systems, since the previous reports primarily studied HUWE1 in *C. elegans* and HEK293T cells [33, 34], while the experiments presented in the current study were conducted in HAP1 cells. Identifying the substrate of HUWE1 that mediates its role as a positive regulator of WNT/CTNNB1 signaling in HAP1 cells should help explain these differences.

We demonstrate that HUWE1 loss reduces WNT signaling in cells containing mutations in some WNT pathway components but not in others (Figs 1-5). These results raise the possibility of targeting the signaling mechanisms by which HUWE1 enhances WNT signaling selectively in tumors harboring mutations in specific WNT pathway components. Eliminating or reducing the activity of HUWE1 itself, which reduces WNT/CTNNB1 signaling in WT HAP1-7TGP, CSNK1A1^KO^ and CTNNB1^ST-A^ cells, is unlikely to be a viable therapeutic strategy due to the pleiotropic effects of HUWE1 on cell physiology, including tumor suppressor functions [39]. However, if the relevant ubiquitylated target of HUWE1 is identified, there may be other ways to phenocopy the effects of HUWE1 loss on WNT signaling more specifically. Phenocopying the effects of HUWE1 loss may not be effective in tumors driven by APC truncations, given that in APC^KO^ cells HUWE1 loss does not reduce WNT signaling due to the role of APC itself in mediating the effects of HUWE1. However, in tumors containing activating mutations in CTNNB1 like those engineered into our CTNNB1^ST-A^ cell line, or mutations in the ZNRF3 or RNF43 tumor suppressors, all of which result in hyperactive WNT signaling in the presence of a functional DC, phenocopying the effects of HUWE1 loss may reduce WNT signaling enough to provide a therapeutic benefit.

We recognize that all the experiments presented in the Results section of this manuscript were conducted in HAP1 cells or derivatives thereof, which could raise concerns about the generality and specificity of our conclusions. We also studied HUWE1 in other cell lines commonly used in WNT signaling research, but our attempts to knock out HUWE1 yielded only partial KOs. We targeted *HUWE1* by CRISPR/Cas9-mediated genome editing in HEK293T-7TG and HEK293T-7TG CSNK1A1^KO^ cells (see Materials and methods). Out of 113 independent clonal cell lines in which we identified mutations in all *HUWE1* alleles, at least one allele had been repaired in frame to encode WT HUWE1 protein (S6A and S6B Figs). This is probably because HUWE1 is a common essential gene as per DEPMAP classification (https://depmap.org/portal/gene/HUWE1?tab=overview), so complete loss of HUWE1 may be lethal in HEK293T cells. However, we have previously shown that microinjection of *HUWE1* mRNA into *Xenopus* embryos results in body axis duplication [15], consistent with a more general role of HUWE1 as a positive regulator of WNT signaling beyond HAP1 cells.

Furthermore, we designed many of our experiments so as to minimize the possibility of non-specific or pleiotropic effects. We knocked out HUWE1 in two independent CSNK1A1^KO^ cell lines with comparable results (Fig 3). We used two different sgRNAs for CRISPR/Cas9-mediated HUWE1 KO in multiple clonal cell lines (Figs 1-3), a different sgRNA for CRISPR base editing of the HUWE1 catalytic residue in multiple CSNK1A1^KO^; HUWE1^C4341R^ clonal cell lines (Fig 5), and another two different sgRNAs for CRISPRi-mediated HUWE1 KD in polyclonal cell populations (Fig 4). In all cases, we found reproducible reductions in WNT/CTNNB1 signaling. We also saw consistent effects of HUWE1 loss in three different genetic backgrounds: WT HAP1-7TGP, CSNK1A1^KO^ and CTNNB1^ST-A^ cells (Figs 1-5). We measured the effects of HUWE1 loss on three or four endogenous WNT target genes and on an 7TGP, an established WNT transcriptional reporter, with comparable outcomes (Figs 1-5). The effects of HUWE1 on WNT signaling could be reversed completely by a relatively short and specific pharmacological treatment with the GSK3A/GSK3B inhibitor CHIR-99021 (Fig 1), and by introducing mutations in some DC components but not others (Figs 3 and 4). Altogether, these results make it very unlikely that the effects of HUWE1 loss are non-specific or due to pleiotropic downregulation of unrelated cellular functions that affect WNT signaling.

Our study also highlights the remarkable potential of HAP1 haploid cells to dissect complex genetic networks in a cell line of human origin [16] through a combination of genome-wide forward genetic screens, loss-of-function and site-directed mutagenesis analyses, and genetic interaction analyses. Despite great advances in CRISPR/Cas-based genome editing technologies during the last decade [40], it remains challenging to knock out two or more alleles of multiple genes and to introduce targeted homozygous point mutations at scale in diploid primary cells, stem cells, and polyploid immortalized cell lines. We could readily do both in HAP1 cells because they have a single allele of most genes. This enabled us to conduct loss-of-function genetic analyses in multiple genetic backgrounds by comparing several HUWE1 KO and control WT clonal cell lines to obtain highly quantitative phenotypic data that confirmed and extended the results of our initial genetic screens (Fig 3). Using CRISPR/Cas9-mediated HDR, we generated a CTNNB1 variant in which we mutated three key phosphorylation sites in the phosphodegron at the single endogenous *CTNNB1* locus, and in a second round of CRISPR/Cas9-mediated genome editing we generated multiple HUWE1 KO and WT cell lines to demonstrate that regulation of WNT signaling by HUWE1 has a component that is independent of CTNNB1 stability (Fig 2). Using CRISPR-mediated base editing, we generated three clonal cell lines containing a point mutation in the catalytic residue of HUWE1 at the single endogenous *HUWE1* locus, which enabled us to demonstrate that the ubiquitin ligase activity of HUWE1 is required for its function in WNT signaling (Fig 5). Finally, we generated single, double, and triple KO mutants for all components of the DC, alone and in certain combinations (11 distinct mutant genetic backgrounds in total) (Table 1 and S4 Fig). Combined with a CRISPRi strategy, this enabled us to carry out an extensive genetic interaction analysis and demonstrate that positive regulation of WNT signaling by HUWE1 is mediated by a subset of DC components (Fig 4 and Table 1). These kinds of genetic analyses would have been practically impossible to conduct in any diploid or polyploid human cell line. We hope this study will inspire other researchers to take advantage of haploid human cell lines, of which there are now many available [41, 42], to unravel other signaling pathways or biological processes in similar ways.

HUWE1 has emerged as an important ubiquitin ligase with many cellular functions [17–20]. Here we show another role for HUWE1 regulating WNT/CTNNB1 signaling. Regulation of CTNNB1 abundance by the DC is the central step in WNT/CTNNB1 signaling. Our discovery that HUWE1 enhances WNT signaling by antagonizing DC-dependent CTNNB1 phosphorylation, thereby increasing CTNNB1 abundance, demonstrates that this crucial step in WNT/CTNNB1 signaling is subject to more nuanced regulation than previously thought. The second mechanism by which HUWE1 enhances WNT signaling independently of CTNNB1 stability is an intriguing additional layer of regulation that remains to be elucidated. Both mechanisms provide new insights into WNT signaling and ubiquitin biology, bridging two research fields that already have many intimate connections.

## Materials and methods

The following Materials and methods relevant to this manuscript have been described previously [15]: cell lines and growth conditions, preparation of WNT3A conditioned media and construction of the HAP1-7TGP WNT reporter haploid cell line.

### Tissue culture media

Complete growth medium (CGM) 1 contains Dulbecco’s Modified Eagles Medium (DMEM) with High Glucose, without L-Glutamine and Sodium Pyruvate (GE Healthcare Life Sciences Cat. # SH30081.01); 1X GlutaMAX-I (Thermo Fisher Scientific Cat. # 35050079); 1X MEM Non-Essential Amino Acids (Thermo Fisher Scientific Cat. # 11140050); 1 mM Sodium Pyruvate (Thermo Fisher Scientific Cat. # 11360070); 40 Units/ml Penicillin, 40 mg/ml Streptomycin (Thermo Fisher Scientific Cat. # 15140122); 10% Fetal Bovine Serum (FBS).

CGM 2 contains Iscove’s Modified Dulbecco’s Medium (IMDM) with L-glutamine, with HEPES, without Alpha-Thioglycerol (GE Healthcare Life Sciences Cat. # SH30228.01); 1X GlutaMAX-I; 40 Units/ml Penicillin, 40 mg/ml Streptomycin; 10% FBS.

### Plasmids

pX330-U6-Chimeric_BB-CBh-hSpCas9 (pX330) (Addgene plasmid # 42230) was a gift from Feng Zhang; pCMV_ABEmax_P2A_GFP (Addgene plasmid # 112101) was a gift from David Liu; MLM3636 (Addgene plasmid # 43860) was a gift from Keith Joung; Lenti-(BB)-EF1a-KRAB-dCas9-P2A-BlastR (Addgene plasmid # 118154) was a gift from Jorge Ferrer; LentiCRISPRv2-mCherry (Addgene plasmid # 99154) was a gift from Agata Smogorzewska; pMDLg/pRRE (Addgene plasmid # 12251), pRSV-Rev (Addgene plasmid # 12253) and pMD2.G (Addgene plasmid # 12259) were a gift from Didier Trono; pCS2-YFP was a gift from Henry Ho; pmCherry was a gift from Jan Carette; pX458-mCherry was generated as described previously [43].

The following plasmids were purchased: pLenti6.2/V5-DEST (Thermo Fisher Scientific Cat. # V36820); pENTR2B (Thermo Fisher Scientific Cat. # A10463); MGC Human AXIN1 Sequence-verified cDNA (Clone ID 5809104) (Horizon Cat. # MHS6278-202833071).

To generate pCMV_ABEmax_P2A_mCherry, mCherry was amplified by PCR from plasmid pmCherry using primers pCMV_ABEmax_P2A_mCherry_Fw (5’-GAA GCA GGC TGG AGA CGT GGA GGA GAA CCC TGG ACC TAT GGT GAG CAA GGG CGA GGA-3’) and pCMV_ABEmax_P2A_mCherry_Rv (5’-CAG ACT TGT ACA GCT CGT CCA TGC CG-3’), designed to include BsmBI and BsrGI restriction sites, respectively. The PCR product was digested with BsmBI and BsrGI and ligated into pCMV_ABEmax_P2A_GFP digested with the same enzymes to replace GFP with mCherry.

To generate pLenti6.2-V5-EXP-N-TERM-S-FLAG-N-hAXIN1, human AXIN1 was amplified by PCR from MGC Human AXIN1 Sequence-verified cDNA (Clone ID 5809104) using primers pENTR2B_SalI_S-FLAG-N_hAXIN1_pcr_fw (5’-GCG CCG GAA CCA ATT CAG TCG ACC CTG CAG GAT GGA TTA CAA GGA CGA CGA TGA CAA GGG CGG CCG CAT GAA TAT CCA AGA GCA GGG TTT CCC CTT GGA CC-3’), containing an N-terminal SalI restriction site followed by a FLAG tag sequence flanked by SbfI and NotI restriction sites, and pENTR2B_XhoI_hAXIN1_pcr_rv (5’-AAA GCT GGG TCT AGA TAT CTC GAG TCA GTC CAC CTT CTC CAC TTT GCC GAT GA-3’), containing a C-terminal XhoI restriction site. The product was digested with SalI and XhoI, and subcloned into pENTR2B digested with the same enzymes. One clone was sequenced completely and subcloned into pLenti6.2/V5-DEST using the Gateway LR Clonase II Enzyme mix.

All constructs were confirmed by sequencing.

### Antibodies

Primary antibodies: purified mouse anti-β-catenin (Clone 14/Beta-Catenin) (1:1000, BD Biosciences Cat. # 610154), rabbit mAb anti-non-phospho (active) β-catenin (Ser33-37-Thr41) (D13A1) (1:1000, Cell Signaling Technology Cat. # 8814), mouse anti-GAPDH (1:4000, Santa Cruz Biotechnology, Cat. # sc-47724), recombinant rabbit anti-Sodium Potassium (Na^+^/K^+^) ATPase [EP1845Y] (1:4000, Abcam Cat. # ab76020), rabbit anti-Lasu1/Ureb1 (HUWE1) (1:1000, Bethyl Laboratories Cat. # A300-486A), rabbit mAb anti-AXIN1 (C76H11) (1:1000, Cell Signaling Technology Cat. # 2087), rabbit mAb anti-AXIN2 (76G6) (1:500, Cell Signaling Technology Cat. # 2151), rabbit mAb anti-GSK-3α/β (D75D3) (1:2000, Cell Signaling Technology Cat. # 5676), mouse anti-APC (NT, clone Ali 12.28) (1:1000, Millipore Sigma, Cat. # MAB3785), rabbit anti-APC (1:1000, Biorbyt Cat. # orb213564), mouse anti-CSNK1A1 (1:250, Santa Cruz Biotechnology, Cat. # sc-74582).

Secondary antibodies: IRDye 800CW donkey anti-mouse IgG (H+L) (1:10000, Li-Cor Cat. # 926-32212), IRDye 680RD donkey anti-rabbit IgG (H+L) (1:10000, Li-Cor Cat. # 925-68073), peroxidase AffiniPure donkey anti-goat IgG (H+L) (1:5000, Jackson ImmunoResearch Laboratories Cat. # 705-035-003), peroxidase AffiniPure goat anti-rabbit IgG (H+L) (1:10000, Jackson ImmunoResearch Laboratories Cat. # 111-035-003), peroxidase AffiniPure donkey anti-mouse IgG (H+L) (1:5000, Jackson ImmunoResearch Laboratories Cat. # 715-035-150), goat anti-mouse IgG (H+L) HRP conjugate (1:10000, Bio-Rad Cat. # 1706516).

Primary and secondary antibodies used for detection with the Li-Cor Odyssey imaging system were diluted in a 1 to 1 mixture of Odyssey Intercept Blocking Buffer (Li-Cor Cat. # 927–40000) and TBST (Tris buffered saline (TBS) + 0.1% Tween-20), and those used for detection by chemiluminescence were diluted in TBST + 5% skim milk. All primary antibody incubations were done overnight at 4°C, and secondary antibody incubations were done for 1 hr at room temperature (RT).

### Construction of mutant HAP1-7TGP cell lines by CRISPR/Cas9-mediated genome editing

Oligonucleotides encoding single guide RNAs (sgRNAs) (S2 File) were selected from a published library [44], or designed using either of two online CRISPR design tools [45, 46] and cloned into either pX330 or pX458-mCherry according to a published protocol [47].

Clonal HAP1-7TGP cell lines were established by transient transfection with either pX330 or pX458-mCherry containing the sgRNA followed by single cell sorting as follows. A transfection mix was prepared by diluting 450 ng of pX330 and 50 ng of pmCherry (used as a cotransfection marker for FACS sorting) or 500 ng of pX458-mCherry in 48 µl Opti-MEM I, adding 2 µl of X-tremeGENE HP and incubating for 20 min at RT. HAP1-7TGP cells or derivatives thereof were reverse-transfected in a well of a 24-well plate by overlaying 0.5 ml of CGM 2 (without antibiotics) containing 6 x 10^5^ cells over the 50 µl of transfection mix. Cells were passaged to a 10 cm dish ∼24 hr post-transfection, using 150 µl of Trypsin-EDTA (0.25%) (Thermo Fisher Scientific Cat. # 25200056) to detach them (reverse-transfection of HAP1 cells caused unusually high adherence, hence the higher trypsin concentration). Three to four days post-transfection, single transfected (mCherry^+^) cells were sorted into 96-well plates containing 200 µl of CGM 2 per well and grown undisturbed for 16 to 18 days. Single colonies were passaged to 24-well plates, and a small number of cells was reserved for genotyping.

For genotyping, genomic DNA was extracted by adding 4 volumes of QuickExtract DNA Extraction Solution (Epicentre Cat. # QE09050) to the cells. Extracts were incubated 10 min at 65°C, 3 min at 98°C, and 5 µl were used as input for PCR amplification of the genomic locus containing the sgRNA target site in 15 µl reactions containing 1X LongAmp Taq reaction buffer, 300 mM of each dNTP, 400 nM of each of the flanking primers indicated in S2 File (most of them designed using the Primer-BLAST online tool from the NCBI) and 0.1 units/µl of LongAmp Taq DNA polymerase (NEB Cat. # M0323L). Amplification of the genomic locus containing the sgRNA target site was confirmed by analysis of the PCR products on a 1% agarose gel and the presence of desired mutations was confirmed by sequencing the amplicons using the primers indicated in S2 File. Given that most engineered cell lines remained haploid, sequencing results were usually unequivocal. Sequencing results for all the clonal cell lines used in the study is presented in S1 File, and for selected clonal cell lines, immunoblot analysis confirmed the absence of the protein products.

Whenever possible, multiple independent mutant cells lines, often generated using two different sgRNAs (see S1 File), were expanded and used for further characterization. For some of the comparisons between WT and mutant cells, multiple individual cell lines confirmed by sequencing to be WT at the sgRNA target site were also expanded and used as controls. To generate double and triple mutant cell lines, a single clonal cell line with the first desired mutation was used in a subsequent round of transfection with either pX330 or pX458-mCherry containing the second and, if applicable, third sgRNAs. Alternatively, WT HAP1-7TGP cells were directly transfected with a combination of pX330 or pX458-mCherry constructs targeting two genes simultaneously.

### Construction of CTNNB1^ST-A^ cell line by CRISPR/Cas9-mediated HDR

Oligonucleotides encoding sgRNAs complementary to exon 3 of *CTNNB1* (S2 File) were designed using either of two online CRISPR design tools [45, 46] and cloned into pX458-mCherry using a published protocol [47].

Clonal CTNNB1^ST-A^ cell lines were established by transient transfection of HAP1-7TGP cells with pX458-mCherry containing the sgRNA, and a single stranded oligonucleotide (ssODN) donor template encoding the desired mutations, called CTNNB1 (ST-A mutant) donor (5’-ATT TGA TGG AGT TGG ACA TGG CCA TGG AAC CAG ACA GAA AAG CGG CTG TTA GTC ACT GGC AGC AAC AGT CTT ACC TGG ACG CTG GAA TCC ATG CTG GTG CCA CTG CCA CAG CTC CTG CTC TGA GTG GTA AAG GCA ATC CTG AGG AAG AGG ATG TGG ATA CCT CCC AAG TCC TGT ATG AGT GGG AAC AGG GAT TTT CTC AG-3’). A transfection mix was prepared by diluting 500 ng pX458-mCherry-CTNNB1-Ex3-sgRNA and 500 ng (8 pmol) ssODN in 48 µl Opti-MEM I. 2 µl of X-tremeGENE HP were added, and the mix was vortexed and incubated for 20 min at RT. The 50 µl mix was placed in an empty well of a 24-well plate and 0.5 ml of CGM 2 containing 6 x 10^5^ cells was seeded onto the mix. The cells were passaged the following day to a 10 cm dish and grown for 3 additional days. Single cells exhibiting high EGFP fluorescence from the 7TGP WNT reporter, presumably due to successful mutagenesis of the CTNNB1 phosphodegron, were sorted, expanded, and genotyped as described above. A single clonal cell line containing point mutations in three of the four targeted sites in the phosphodegron (S2A Fig and S1 File) was used for all subsequent experiments.

### Construction of HUWE1 catalytic mutant CSNK1A1^KO^; HUWE1^C4341R^ cell lines by base editing

An oligonucleotide encoding an sgRNA complementary to exon 83 of *HUWE1* (S2 File) was designed to include the targeted nucleotide within the editing window of the base editor ABEmax (positions 4-8 in the protospacer) using BE-Hive (https://www.crisprbehive.design), an online base editing sgRNA design tool [48], and cloned into MLM3636 according to a published protocol (Joung Lab gRNA cloning protocol: https://media.addgene.org/data/plasmids/43/43860/43860-attachment_T35tt6ebKxov.pdf). A transfection mix was prepared by diluting 750 ng pCMV-ABEmax-P2A-mCherry and 250 ng MLM3636-HUWE1-C4341R-sgRNA1 in 50 µl Opti-MEM I, adding 2 µl of X-tremeGENE HP and incubating for 20 min at RT. CSNK1A1^KO^ cells were reverse-transfected in a well of a 24-well plate by overlaying 0.5 ml of CGM 2 (without antibiotics) containing 6 x 10^5^ cells over the 50 µl of transfection mix. Cells were passaged to a 6 cm dish ∼24 hr post-transfection, using 150 µl of Trypsin-EDTA (0.25%) (Thermo Fisher Scientific Cat. # 25200056) to detach them. Three days post-transfection, single transfected (mCherry^+^) cells were sorted into 96-well plates containing 200 µl of CGM 2 per well and grown undisturbed for 16 to 18 days. Cells were expanded and genotyped as described above.

### Targeting *HUWE1* by CRISPR/Cas9 in HEK293T-7TG and HEK293T-7TG CSNK1A1^KO^ cells

Oligonucleotides HUWE1-IVT-2503-F and HUWE1-IVT-2503-R encoding sgRNAs complementary to exon 6 of *HUWE1* (S2 File) were designed using sgRNA Scorer 2.0 [49] and cloned into LentiCRISPRv2-mCherry previously digested with BsmBI. HEK293T-7TG is a clonal cell line derived from HEK293T cells that contains a fluorescent WNT reporter. HEK293T-7TG CSNK1A1^KO^ is a clonal cell line derived from HEK293T-7TG cells in which CSNK1A1 has been knocked out. Construction of both cell lines will be described elsewhere.

Clonal HEK293T-7TG and HEK293T-7TG CSNK1A1^KO^ cell lines in which HUWE1 was targeted by CRISPR/Cas9 were established by transient transfection with LentiCRISPRv2-mCherry containing the sgRNAs followed by single cell sorting. ∼24 hr before transfection, 8 x 10^4^ HEK293T-7TG or HEK293T-7TG CSNK1A1^KO^ cells per well were seeded in 24-well plates and grown in CGM 1. On the day of transfection, CGM 1 was replaced with 450 µl of antibiotic-free CGM 1. 50 µl of a transfection mixture containing 500 ng LentiCRISPRv2-mCherry and 1 µl of X-tremeGENE^™^ HP DNA Transfection Reagent (Millipore Sigma, Cat # 06366236001) prepared in OptiMEM were added dropwise. ∼24 hr post-transfection, cells were transferred to a 6 cm dish, and ∼72 hr post-transfection, single transfected (mCherry^+^) cells were sorted into 96-well plates containing 200 µl of CGM 1 media per well and grown undisturbed for 16 days. Single colonies were expanded by passaging to 24-well plates, and 10 μl of cell suspension were reserved for genotyping.

For genotyping, genomic DNA was extracted by adding 4 volumes of QuickExtract DNA Extraction Solution (Epicentre, Cat # QE09050) to the cells. Extracts were incubated for 10 min at 65°C, 3 min at 98°C, and 5 µl were used as input for PCR amplification of the *HUWE1* target site in 15 µl reactions containing 1X LongAmp Taq reaction buffer, 300 mM of each dNTP, 400 nM of each of the flanking primers PS1057-NGS-F and PS1057-NGS-R (S2 File) and 0.1 units/µl of LongAmp Taq DNA polymerase (NEB Cat. # M0323L). In a second amplification step, complete Illumina adapter sequences (F: 5’-AAT GAT ACG GCG ACC ACC GAG ATC TAC AC <8 bp barcode> AC ACT CTT TCC CTA CAC GAC GCT CTT CCG ATC* T-3’ ; R: 5’-CAA GCA GAA GAC GGC ATA CGA GAT <8 bp barcode> G TGA CTG GAG TTC AGA CGT GTG CTC TTC CGA TC*T-3’; * indicates a phosphorothioate (PTO) linked base) were added and the amplicons were sequenced on the MiSeq system (Illumina). FASTQ sequencing files were analyzed using the branch 1.1 version [50] of a previously described analysis pipeline (https://github.com/rajchari2/ngs_amplicon_analysis). Total (dark blue) and out-of-frame (light blue) mutation rates were calculated and plotted (S6 Fig).

### Preparation of lentivirus, lentiviral transduction, and selection of HUWE1 KD and AXIN1-overexpressing polyclonal cell populations

The transfer plasmid used to generate HUWE1 KD cell lines by CRISPRi was Lenti-(BB)-EF1a-KRAB-dCas9-P2A-BlastR. The transfer plasmid used to generate cell lines overexpressing AXIN1 was pLenti6.2-V5-EXP-N-TERM-S-FLAG-N-hAXIN1. ∼24 hr before transfection, 21 x 10^6^ HEK293T cells were plated in 20 ml of CGM 1 without antibiotics in a T-175 flask. A transfection mixture was prepared by diluting 9.3 µg of transfer plasmid, 7 µg of pMDLg/pRRE, 7 µg of pRSV-Rev, 4.66 µg of pMD2.G, 1.05 µg pCS2-YFP (as a cotransfection marker), and 87.15 µl of 1 mg/ml polyethylenimine (PEI) in a final volume of 1 ml serum-free DMEM. The mixture was incubated for 20 min at RT and added to the culture media in the flasks. The day after transfection, the media was replaced with 18 ml of CGM1 containing a total of 20% FBS without antibiotics. ∼48 hr after transfection, the media was collected (first viral harvest), centrifuged at 1000 x g for 5 min to remove cell debris, and the supernatant was reserved at 4°C. 18 ml of fresh media were added to the flask of cells. ∼72 hr after transfection, the media was collected (second viral harvest), centrifuged as before, and the supernatant was pooled with the first viral harvest. The pooled supernatant was filtered through 0.45 µm filters (Acrodisc syringe filters with 0.45 µm Supor membrane, Pall Corporation Cat. # 4654). The filtered media containing lentiviral particles was aliquoted, snap frozen in liquid nitrogen, and stored at -80°C.

For smaller scale preparations of the lentivirus used for AXIN1 overexpression, the above protocol was followed but the lentivirus was prepared using 293FT cells in in T-25 flasks and all quantities and volumes were scaled down by ∼1/7. 3 x 10^6^ 293FT cells were plated in 5 ml of CGM 1 without antibiotics in a T-25 flask. A transfection mixture was prepared by diluting 1.33 µg of transfer plasmid, 1 µg of pMDLg/pRRE, 1 µg of pRSV-Rev, 0.66 µg of pMD2.G, 0.15 µg pCS2-YFP (as a cotransfection marker), and 69.4 µg/ml PEI in a final volume of 180 µL serum-free DMEM. The mixture was incubated for 20 min at RT and added to the culture media in the flasks. The day after transfection, the media was replaced with 2.5 ml of CGM 1 containing 20% FBS without antibiotics and the viral supernatants were collected and processed as described above.

Approximately 24 hr before transduction, 2.5 x 10^5^ HAP1-7TGP cells or derivatives thereof were seeded in a 6-well plate. Cells were transduced by adding 1 ml of lentivirus-containing supernatant mixed with 1 ml of CGM 2 and 4.4 µg/ml polybrene. ∼24 hr post-transduction, cells were passaged to 10 cm dishes and selected with 8 µg/ml blasticidin in CGM 2 for ∼96 hr. Untransduced cells from each genetic background were treated in parallel with 8 µg/mL blasticidin to ensure that all cells were killed by the time selection of transduced cells was complete.

### Analysis of WNT reporter fluorescence

To measure WNT reporter activity in HAP1-7TGP cells or derivatives thereof, ∼24 hr before treatment cells were seeded in 24-well plates at a density of 8 x 10^4^ per well and grown in 0.5 ml of CGM 2. Cells were treated for 24 hr with the indicated concentrations of WNT3A CM diluted in CGM 2. Cells were washed with 0.5 ml PBS, harvested in 150 µl of Trypsin-EDTA (0.05%) (Thermo Fisher Scientific Cat. # 25300054), resuspended in 450 µl of CGM 2, and EGFP fluorescence was measured by FACS on either a SA3800 Spectral Cell Analyzer (Sony Biotechnology) or a CytoFLEX S Flow Cytometer (Beckman Coulter). Typically, fluorescence data for 5,000–50,000 singlet-gated cells was collected and, unless indicated otherwise, the median EGFP fluorescence ± standard error of the median (SEM = 1.253 s/n, where s = standard deviation and n = sample size) was used to represent the data.

To measure WNT reporter activity in cells treated with the GSK3A/B inhibitor CHIR-99021, ∼24 hr before treatment cells were seeded in 6-well plates at a density of 0.5 x 10^6^ per dish. Cells were treated the following day with 10 µM CHIR-99021 (CT99021) (Selleckchem Cat. # S2924) or an equivalent volume of DMSO vehicle diluted in CGM 2 for 48 hr, replacing the media with fresh CHIR-99021 or DMSO in CGM 2 after 24 hr of treatment. Cells were washed with 2 ml PBS, harvested in 0.5 ml of 0.05% Trypsin-EDTA, resuspended in 1.5 ml of CGM 2, and EGFP fluorescence was measured as described above.

### Quantitative (q)RT-PCR analysis

Approximately 24 hr before treatment, cells were seeded in 24-well plates at a density of 3 x 10^5^ per well and grown in 0.5 ml of CGM 2. Cells were treated for 24 hr with 50% WNT3A CM in CGM 2 where indicated. The medium was removed, cells were washed once with PBS and harvested in 400 µl of TRIzol Reagent (Thermo Fisher Scientific Cat. # 15596018). Extracts were processed according to the manufacturer’s protocol, taking the appropriate precautions to avoid contamination with nucleases, and total RNA was resuspended in 20 µl of DEPC-treated water (Thermo Fisher Scientific Cat. # AM9920). To synthesize cDNA, 125 ng of RNA were diluted in 2 µl DEPC-treated water and incubated with 0.25 µl 10X ezDNase buffer and 0.25 µl ezDNase enzyme for 5 min at 37°C to digest DNA contaminants. After DNase treatment, 1 µl of DEPC-treated water and 1 µl of SuperScript^TM^ IV VILO^TM^ MM (Invitrogen Cat. # 11766500) were added and the reaction was incubated for 10 min at 25°C, 10 min at 50°C, and 5 min at 85°C. For each primer pair, a cDNA dilution series from a representative sample was analyzed to ensure that target amplification was linear across a sufficiently broad range of cDNA concentrations. cDNA was diluted 1:100 in water, and 5 µl were mixed with 5 µl of Power SYBR Green PCR Master Mix (Applied Biosystems Cat. # 4367659) containing 400 nM each of forward and reverse primer (S2 File). Triplicate reactions for each cDNA and primer pair were prepared in a MicroAmp Optical 384-well Reaction Plate (Thermo Fisher Scientific Cat. # 4309849), sealed with MicroAmp Optical Adhesive Film (Thermo Fisher Scientific Cat. # 4311971) and run using standard parameters in a QuantStudio 5 Real-Time PCR System (Applied Biosystems). Thermo Fisher cloud design and analysis software (DA2) was used to calculate the average relative abundance of *AXIN2*, *RNF43*, *TNFRSF19*, or *NKD1* mRNA normalized to *HPRT1* mRNA, and fold-changes in mRNA abundance were calculated as the quotient between the experimental and reference samples, with appropriate error propagation of the respective standard deviations (SD).

### Immunoblot analysis and quantification of soluble CTNNB1 from membrane-free supernatants (MFS)

Approximately 24 hr before treatment, cells were seeded in 6 cm dishes at a density of 2.5 x 10^6^ per dish and grown in 5 ml of CGM 2. Cells were treated for 24 hr with 50% WNT3A CM in CGM 2 where indicated. Cells were harvested, lysed by hypotonic shock, and extracts were prepared as follows, with all handling done at 4°C. Cells were washed twice with ∼5 ml cold PBS and twice with ∼5 ml cold 10 mM HEPES pH 7.4. Residual buffer was removed, and 100 µl of ice-cold SEAT buffer (10 mM triethanolamine/acetic acid pH 7.6, 250 mM sucrose, 1X SIGMAFAST Protease Inhibitor Cocktail Tablets EDTA-free (Sigma-Aldrich Cat. # S8830), 25 µM MG132 (Sigma-Aldrich Cat. # C2211), 1X PhosSTOP (Roche Cat. # 04906837001), 1 mM NaF, 1 mM Na_3_VO_4_, 1 mM dithiothreitol (DTT), 62.5 U/ml Benzonase Nuclease (EMD Millipore Cat. # 70664), 1 mM MgCl_2_) were added to the cells. Cells were scraped using a cell lifter (Corning Cat. # 3008), transferred to 2-ml centrifuge tubes and disrupted mechanically by triturating 10 times. Crude extracts were centrifuged for 20 min at 20,000 x g to pellet membranes and other insoluble cellular material, and the MFS was carefully removed, avoiding contamination from the pellet. The MFS was flash-frozen in liquid nitrogen and stored at -80°C until further processing.

Extracts were thawed quickly at RT and transferred to ice. The protein concentration in the MFS was quantified with the Pierce BCA Protein Assay Kit (Thermo Fisher Scientific Cat. # 23225), using BSA as a standard, and samples were normalized by dilution with SEAT buffer. The MFS was diluted with 4X LDS sample buffer (Thermo Fisher Scientific Cat. # NP0007) supplemented with 50 mM tris(2-carboxyethyl)phosphine (TCEP), incubated for 45 min at RT or heated at 95°C for 10 min, and 30 µg of total protein were electrophoresed alongside Precision Plus Protein All Blue Prestained Protein Standards (Bio-Rad Cat. # 1610373) in 4-15% TGX Stain-Free protein gels (BioRad, various Cat. numbers) at 75 V for 15 min and 100 V for 1 hr 15 min using 1X Tris/Glycine/SDS running buffer (BioRad Cat. # 1610772). Following electrophoresis, the gel was briefly activated with UV light using a Chemidoc imager (BioRad) to covalently label proteins in the gel with Stain-Free fluorochromes.

Proteins were transferred to PVDF membranes in a Criterion Blotter apparatus (Bio-Rad Cat. # 1704071) at 60 V for 2 hr using 1X Tris/Glycine transfer buffer (BioRad Cat. # 1610771) containing 20% methanol. Following transfer, the membrane was imaged using the ChemiDoc imager, and the total protein in each lane was quantified. Membranes were cut, blocked with Odyssey Blocking Buffer (Li-Cor Cat. # 927–40000), incubated with mouse anti-β-catenin or mouse anti-GAPDH primary antibodies, washed with TBST, incubated with IRDye 800CW donkey anti-mouse IgG secondary antibody, washed with TBST followed by TBS, and imaged using a Li-Cor Odyssey imaging system. Acquisition parameters in the manufacturer’s Li-Cor Odyssey Image Studio Lite software were set so as to avoid saturated pixels in the bands of interest, and bands were quantified using background subtraction. The integrated intensity for CTNNB1 was normalized to the total protein (or in some cases to the average of total protein and the integrated intensity for GAPDH) in the corresponding lane. The average ± SD normalized CTNNB1 intensity from duplicate blots was used to represent the data.

For CHIR-99021 or DMSO vehicle treated cells, ∼24 hr before treatment cells were seeded in 6 cm dishes at a density of 2 x 10^6^ per dish and treated the following day with 10 µM CHIR-99021 or DMSO for 48 hr. The media was replaced with fresh CHIR-99021 or DMSO in CGM 2 after 24 hr of treatment.

### Immunoblot analyses of soluble HUWE1, APC and CSNK1A1 from MFS

Some of the same membranes used to blot for soluble CTNNB1 were cut and used to blot for other proteins as indicated in the same figures. Blots were incubated with rabbit anti-Lasu1/Ureb1 (HUWE1), mouse anti-APC and mouse anti-CSNK1A1 primary antibodies. The following secondary antibodies were used: for HUWE1, IRDye 680RD donkey anti-rabbit IgG, for APC, IRDye 800CW donkey anti-mouse IgG (both were imaged using a Li-Cor Odyssey imaging system) and for CSNK1A1, peroxidase AffiniPure goat anti-mouse IgG (developed using SuperSignal West Femto Maximum Sensitivity Substrate (Thermo Fisher Scientific Cat. # 34095)).

### Immunoblot analysis of total HUWE1, APC, CTNNB1, AXIN1, AXIN2, GSK3A/B and CSNK1A1 from whole cell extracts (WCE)

Approximately 72 hr before harvest, cells were seeded in 10 cm dishes at a density of 3 x 10^6^ per dish and grown in 10 ml of CGM 2. Cells were harvested in 2 ml Trypsin-EDTA (0.05%) and resuspended in 6 ml CGM 2. 10 x 10^6^ cells were centrifuged at 400 x g for 5 min, washed in 5 ml PBS, and centrifuged at 400 x g for 5 min. The supernatant was aspirated, and the cell pellets were flash-frozen in liquid nitrogen and stored at -80°C. Pellets were thawed quickly at RT and transferred to ice. All subsequent steps were done on ice. The cell pellets were resuspended in 150 µl of ice-cold RIPA lysis buffer (50 mM Tris-HCl pH 8.0, 150 mM NaCl, 2% NP-40, 0.25% deoxycholate, 0.1% SDS, 1X SIGMAFAST protease inhibitors, 1 mM MgCl_2_, 62.5 U/ml Benzonase Nuclease, 1 mM DTT, 10% glycerol), sonicated in a Bioruptor Pico sonication device (Diagenode) 4 x 30 s in the ultra-high setting, centrifuged 10 min at 20,000 x g and the supernatant (WCE) was recovered.

The protein concentration in the WCE was quantified using the Pierce BCA Protein Assay Kit. Samples were normalized by dilution with RIPA lysis buffer, further diluted with 4X LDS sample buffer supplemented with 50 mM TCEP, incubated for 45 min at RT, and 30 µg of total protein were electrophoresed alongside Precision Plus Protein All Blue Prestained Protein Standards in 4-15% Criterion TGX Stain-Free protein gels at 75 V for 15min, and 100 V for 1 hr and 15 min using 1X Tris/Glycine/SDS running buffer.

Proteins were transferred at 60 V for 2 hr to PVDF membranes using 1X Tris/Glycine transfer buffer containing 20% methanol, and membranes were cut and blocked with either Odyssey Intercept Blocking Buffer or TBST, 5% skim milk. Blots were incubated with rabbit anti-Lasu1/Ureb1 (HUWE1), rabbit anti-APC, mouse anti-β-catenin, rabbit anti-AXIN1, rabbit anti-AXIN2, rabbit anti-GSK3A/B, mouse anti-CSNK1A1 and mouse anti-GAPDH (as a loading control) primary antibodies, washed with TBST, incubated with Peroxidase AffiniPure anti-rabbit or anti-mouse secondary antibodies, washed with TBST followed by TBS, and developed with SuperSignal West Femto.

### Immunoblot analysis and quantification of non-phospho-CTNNB1 (S33/S37/T41) and total CTNNB1 from WCE

Approximately 24 hr before treatment, cells were seeded in 6 cm dishes at a density of 2 x 10^6^ per dish and treated the following day with 10 µM CHIR-99021 or an equivalent volume of DMSO vehicle for 48 hr. The media was replaced with fresh CHIR-99021 or DMSO in CGM 2 after 24 hr of treatment. Cells were harvested in 1 ml Trypsin-EDTA (0.05%) and resuspended in 3 ml CGM 2. Cells were centrifuged at 400 x g for 5 min, washed in 5 ml PBS, and centrifuged at 400 x g for 5 min. The above protocol for immunoblot analysis of total proteins from WCE was followed, except that protein samples were heated at 95°C for 10 min prior to electrophoresis, and the total protein in each lane was quantified as follows and used for normalization. Following electrophoresis, the gel was briefly activated with UV light using a Chemidoc imager (BioRad) to covalently label proteins in the gel with Stain-Free fluorochromes. Following transfer, the membrane was imaged using the ChemiDoc, and the total protein in each lane was quantified. The blots were incubated with rabbit non-phospho (active) β-catenin (Ser33-37-Thr41), mouse anti-β-catenin or mouse anti-GAPDH (as a loading control) primary antibodies, IRDye 680RD donkey anti-rabbit IgG or IRDye 800CW donkey anti-mouse IgG secondary antibodies, and imaged using a Li-Cor Odyssey imaging system.

### Quantitative dot blot of HUWE1 from WCE

3 µl WCE containing 8 µg protein were spotted onto nitrocellulose membrane for each sample in triplicate. The membrane was allowed to dry for 15 min prior to staining with Revert 520 Total Protein Stain (Li-Cor Cat. # 926-10010) according to the manufacturer’s protocol (https://www.licor.com/documents/1o8anlg26tnwqkj135ki6bo61fy4ztmi). The membrane was imaged on the Li-Cor Odyssey M imaging system using the 520 nm channel to obtain a total protein quantification for normalization. The membrane was then blocked with Odyssey Blocking Buffer, incubated with rabbit anti-Lasu1/Ureb1 (HUWE1), washed with TBST, incubated with IRDye 680RD donkey anti-rabbit IgG secondary antibody, washed with TBST followed by TBS, and imaged using the Li-Cor Odyssey M imaging system. Acquisition parameters in the manufacturer’s Li-Cor Odyssey Image Studio Lite software were set so as to avoid saturated pixels in the dots of interest, and dots were quantified using background subtraction. The integrated intensity for HUWE1 was normalized to that for Revert 520 Total Protein Stain in the same blot, and the average ± SD from triplicate dot blots was used to represent the data. The specificity of the HUWE1 signal was confirmed by comparing dot blots of WCE from CSNK1A1^KO^ and CSNK1A1^KO^; HUWE1^KO^ cells. All normalized HUWE1 intensity values were within the linear range of a standard curve prepared from dot blots of a serial dilution of WCE from CSNK1A1^KO^ cells.

### Preparation of figures and statistical analysis

Figures were prepared using PowerPoint (Microsoft). Table 1, S1 and S2 Files were prepared using Excel (Microsoft). Graphs were prepared and statistical analysis performed using Prism 6 (GraphPad) or Excel. Details of the statistical tests used are given in the figure legends. Significance is indicated as ****(p<0.0001), *** (p<0.001), ** (p<0.01), * (p<0.05) or n.s. (not significant). Pictures of immunoblots were adjusted for contrast and brightness only when necessary for clarity using Image Studio Lite (LiCor), and were arranged in PowerPoint.

## Acknowledgments

AML was supported by the Intramural Research Program of the National Institutes of Health, National Cancer Institute, Center for Cancer Research. We thank Caleb K. Sinclear for help with experiments, Lebensohn Lab members and members of the Laboratory of Cellular and Molecular Biology for input on the project, Rajat Rohatgi for feedback on the manuscript, and the Center for Cancer Research Genomics Core and the Laboratory of Genome Integrity Flow Cytometry Core for assistance and services. The authors declare no conflicting financial interests.

---

**S1 Fig.**
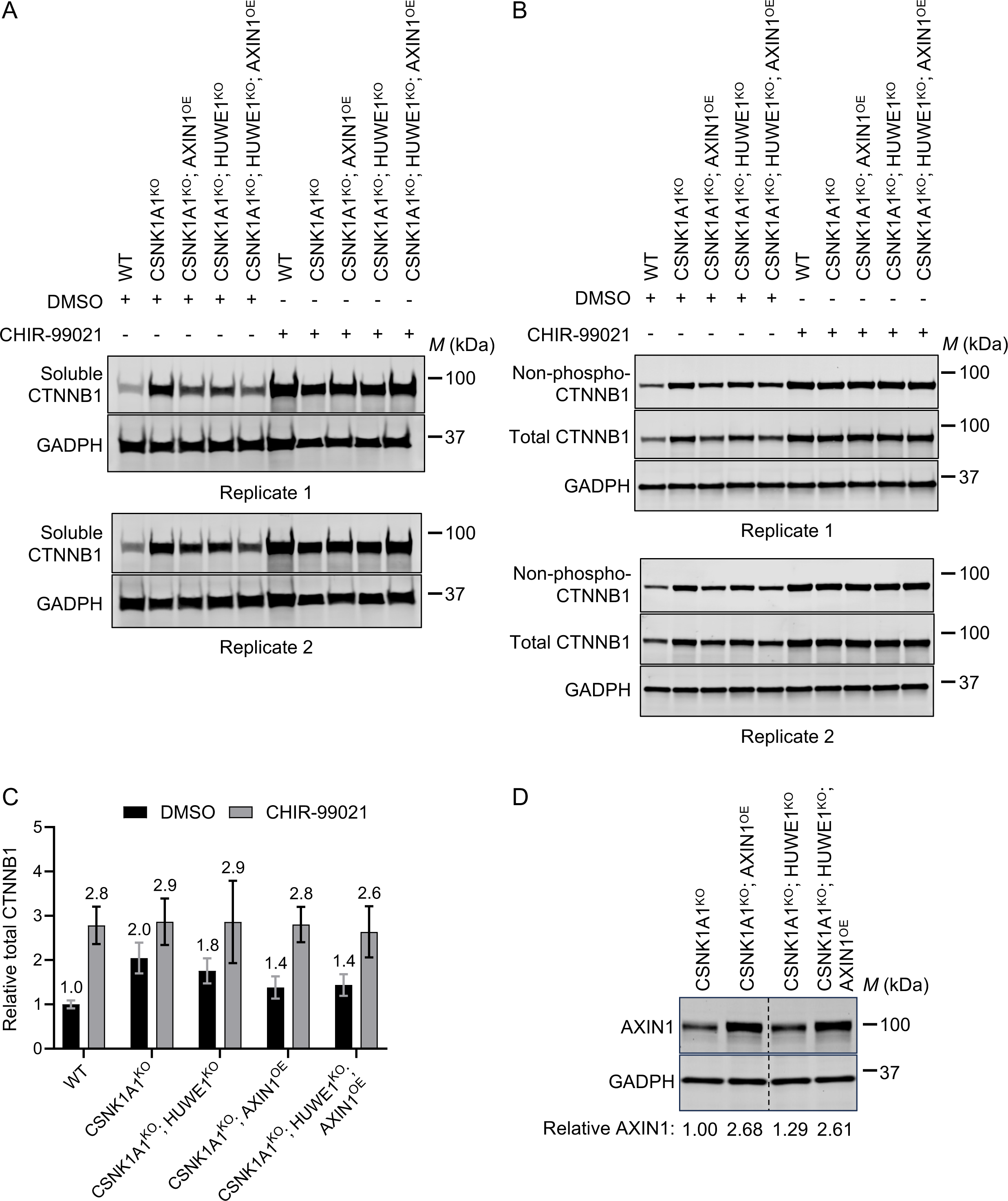
HUWE1 and AXIN1 reciprocally regulate WNT signaling by modulating GSK3A/GSK3B-dependent CTNNB1 phosphorylation and abundance. (A-C) We note that the data for WT HAP-7TGP, CSNK1A1^KO^ and CSNK1A1^KO^; HUWE1^KO^ cells is discussed in the first section of the results, while the data for CSNK1A1^KO^; AXIN1^OE^ and CSNK1A1^KO^; HUWE1^KO^; AXIN1^OE^ cells is discussed in a later section of the results subtitled “HUWE1 enhances WNT signaling by antagonizing the DC.” Cells were treated with DMSO vehicle or 10 µM of the GSK3A/GSK3B inhibitor CHIR-99021 for 48 hr as indicated. (A) Immunoblots of soluble CTNNB1 from MFS, used for quantification in Fig 1B. (B) Immunoblots of non-phospho-CTNNB1 (S33/S37/T41) and total CTNNB1 from WCE, used for quantification in Fig 1C and S1C Fig, respectively. (C) Total CTNNB1 abundance (CTNNB1 intensity normalized to total protein, average ± SD from duplicate immunoblots shown in S1B) in WCE of the indicated cell lines, relative to WT HAP1-7TGP cells treated with DMSO. (D) Immunoblot analysis of total AXIN1 from WCE of the indicated cell lines used in A-C, and in Fig 1. The polyclonal cell populations overexpressing AXIN1 were generated as described in Materials and methods. AXIN1 abundance (AXIN1 intensity normalized to GAPDH intensity), relative to CSNK1A1^KO^ cells, is indicated below the blots.

**S2 Fig.**
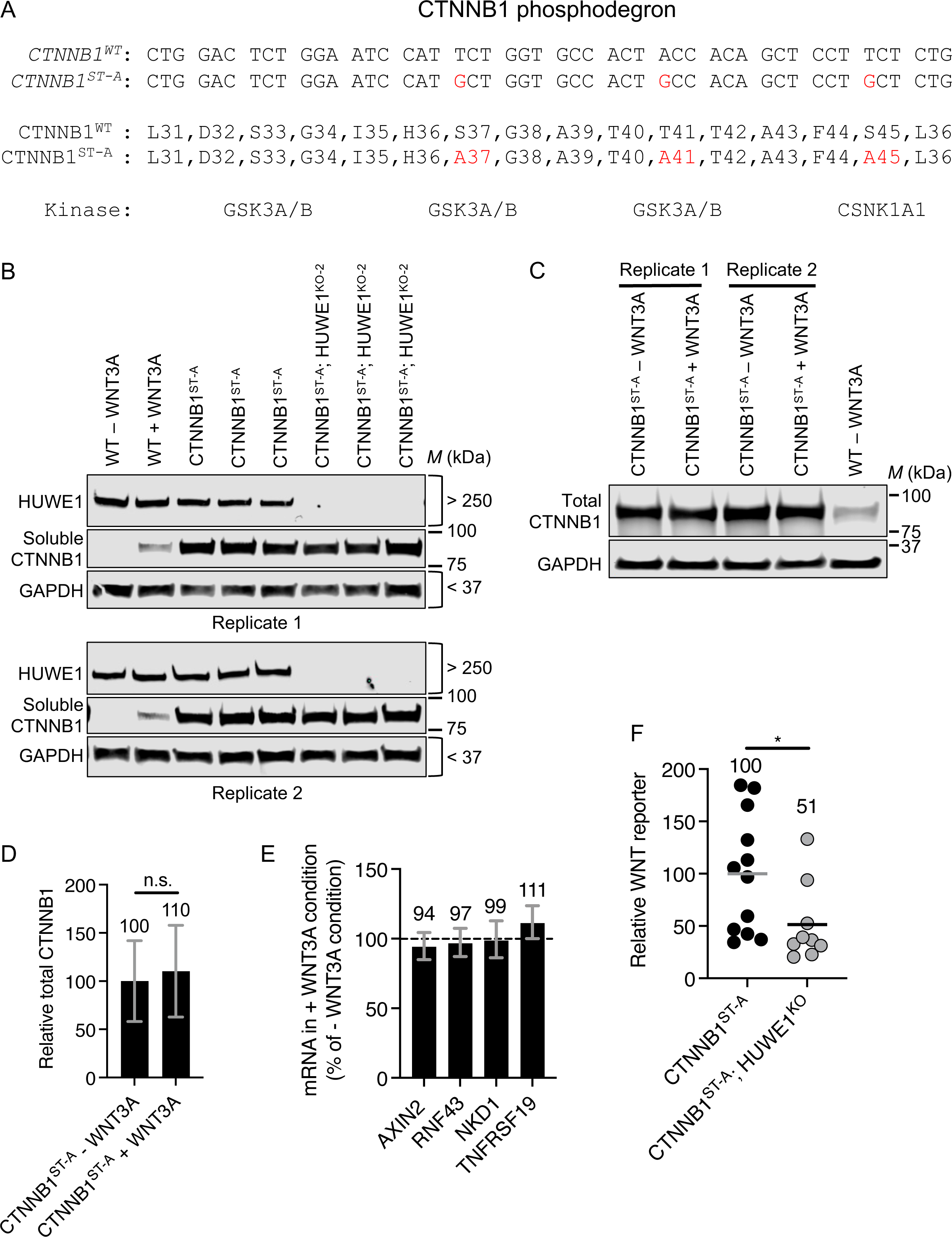
HUWE1 enhances WNT signaling through a mechanism independent of CTNNB1 stability. (A) Genomic nucleotide and corresponding amino acid sequences comprising the CTNNB1 phosphodegron of WT HAP1-7TGP and CTNNB1^ST-A^ cells. The kinases that phosphorylate S or T residues in the phosphodegron are indicated. Nucleotides and amino acids in red indicate mutations. (B) Immunoblots of soluble HUWE1 and CTNNB1 from MFS of the indicated cell lines. The CTNNB1 immunoblots were used for quantification in Fig 2A. (C-E) Treatment of CTNNB1^ST-A^ cells with WNT3A does not promote further accumulation of soluble CTNNB1 and does not further increase WNT target gene expression. Cells were treated with 50% WNT3A CM for 24 hr where indicated. (C) Immunoblots of total CTNNB1 from WCE used for quantification in D. (D) Total CTNNB1 abundance (CTNNB1 intensity normalized to total protein and GAPDH intensity, average ± SD from duplicate lanes of the immunoblots shown in C) in WCE of CTNNB1^ST-A^ cells treated with WNT3A CM, relative to untreated CTNNB1^ST-A^ cells. Significance was determined by unpaired t-test with Welch’s correction. (E) mRNA abundance (average ± SD *AXIN2*, *RNF43*, *TNFRSF19*, or *NKD1* mRNA normalized to *HPRT1* mRNA, each measured in triplicate reactions) in CTNNB1^ST-A^ cells treated with WNT3A CM, reported as percentage of the mRNA abundance in untreated CTNNB1^ST-A^ cells. (F) WNT reporter activity (median EGFP fluorescence from 5,000 singlets) for the indicated cell lines, relative to the average for CTNNB1^ST-A^ cells. Each circle represents a unique clonal cell line (determined by genotyping, S1 File), and the average of 9-12 independent clones for each genotype is indicated by a horizontal line and quantified above each group of circles. Significance was determined by unpaired t-test with Welch’s correction.

**S4 Fig.**
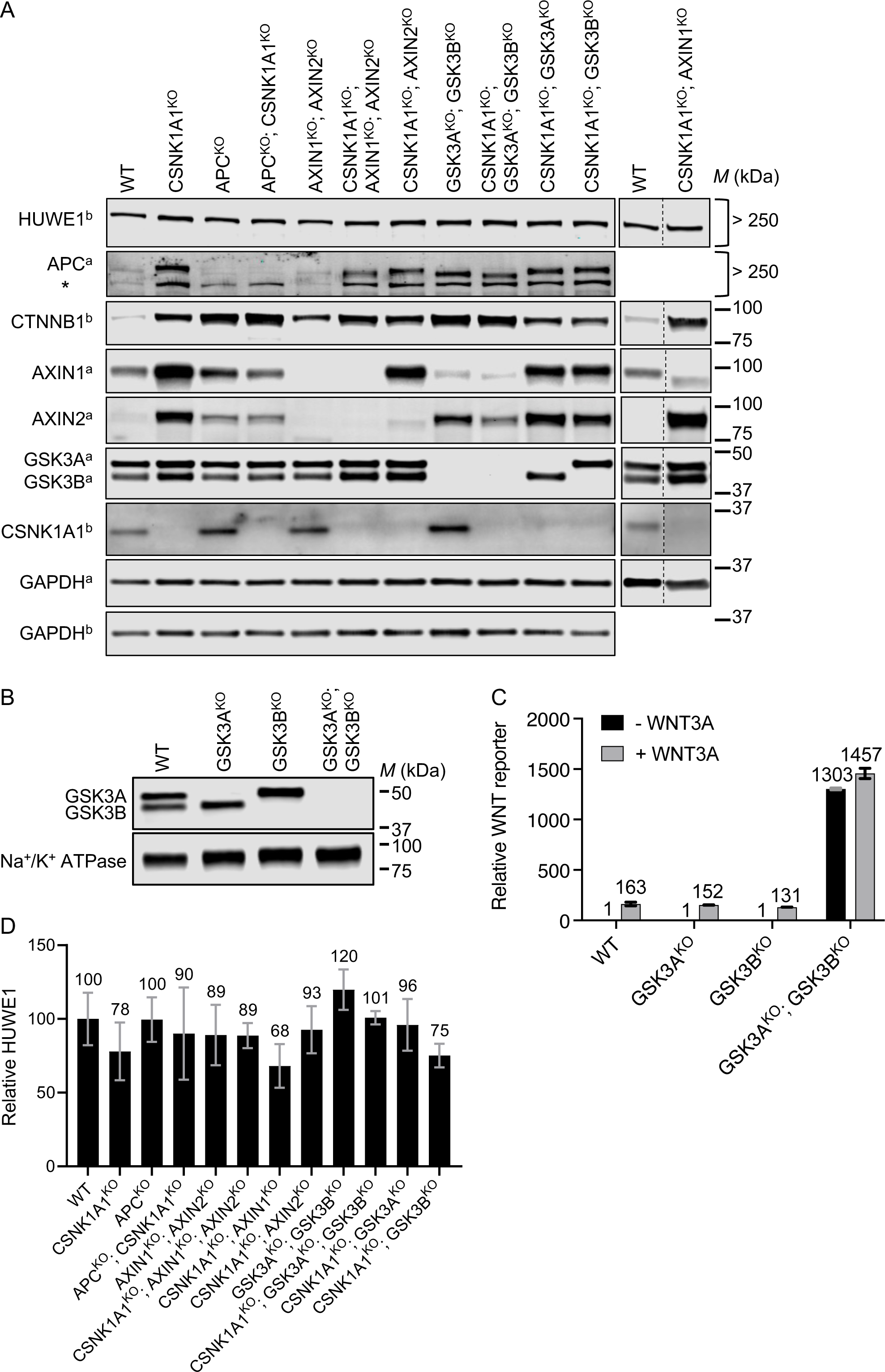
HUWE1 enhances WNT signaling through mechanisms mediated by a subset of DC components including APC, AXIN1 and GSK3A or GSK3B. (A) Immunoblot analysis of total protein from WCE of the indicated clonal cell lines used for CRISPRi-mediated HUWE1 KD in Fig 4 and S5 Fig. The AXIN1 and AXIN2 immunoblots of CSNK1A1^KO^; AXIN1^KO^ and CSNK1A1^KO^; AXIN2^KO^ cells, respectively, exhibited bands of lower abundance and molecular weight than their respective counterparts in WT HAP1-7TGP cells. These bands may represent residual truncated protein products, but in both cases frameshift mutations in the single allele of the respective genes (S1 File) predicted the absence of full-length, WT proteins. * indicates a non-specific band observed with the rabbit anti-APC antibody. The “a” and “b” superscripts next to the protein names indicate which of two membranes the corresponding strips were cut from. Dashed vertical lines indicate a rearrangement of samples within the same blot. (B, C) GSK3A and GSK3B are functionally redundant in WNT signaling in HAP1 cells. The same cell lines were used in B and C. (B) Immunoblot analysis of total GSK3A and GSK3B from WCE of the indicated cell lines. (C) WNT reporter activity (median EGFP fluorescence from 50,000 singlets was measured for biological duplicates of a single clone, and the average ± SD of the two measurements was calculated) relative to untreated WT HAP1-7TGP cells. (D) HUWE1 abundance, quantified by dot blots, in the clonal cell lines used for CRISPRi-mediated HUWE1 KD in Fig 4 and S5 Fig. Total HUWE1 abundance (HUWE1 intensity normalized to total protein, average ± SD from triplicate dot blots) in WCE of the indicated cell lines, relative to WT HAP1-7TGP cells. Significance was determined by unpaired t-test with Welch’s correction, and the difference in HUWE1 abundance between each mutant cell line and WT HAP1-7TGP cells was not significant (not depicted).

**S5 Fig.**
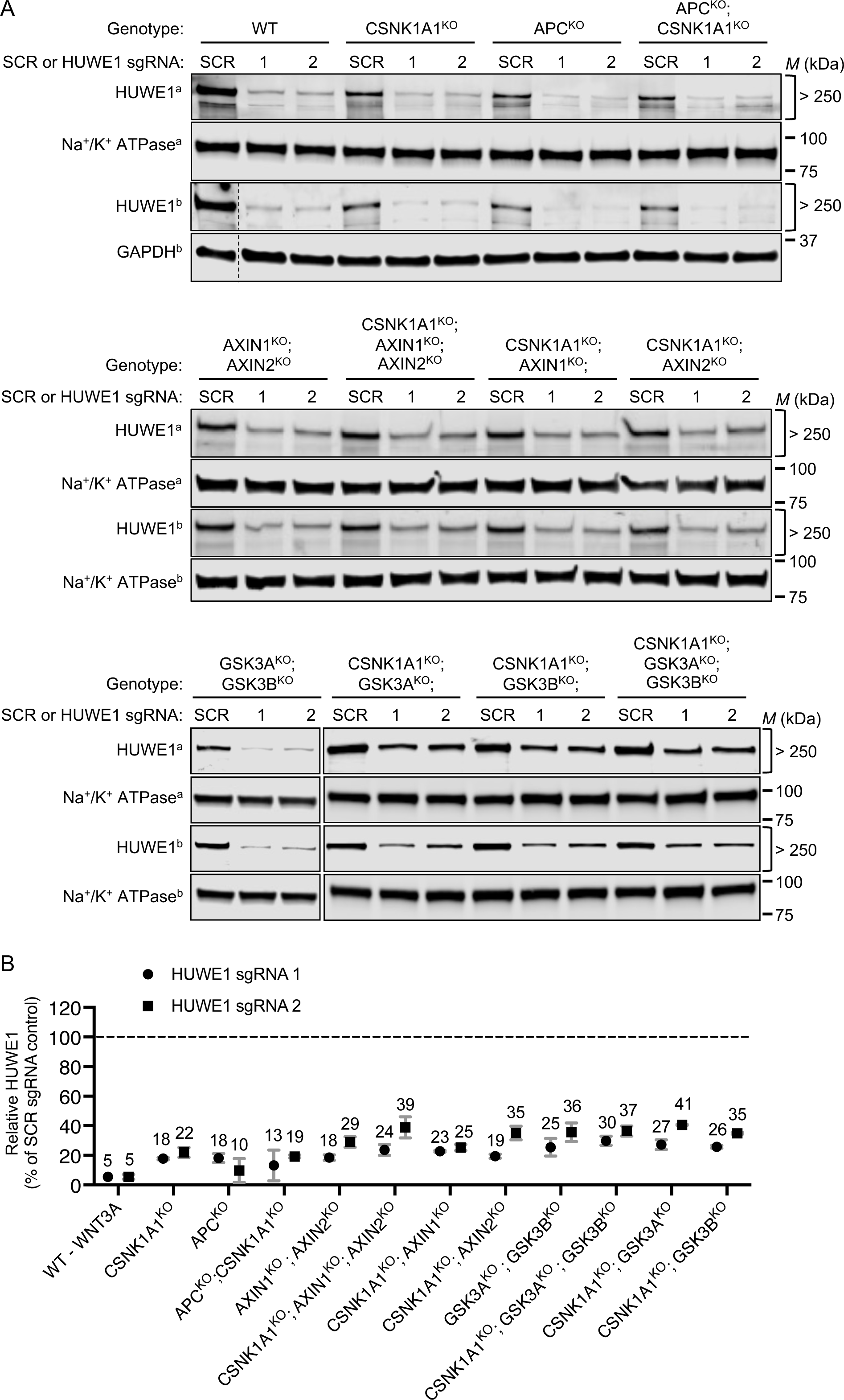
Quantification of CRISPRi-mediated HUWE1 KD in various genetic backgrounds. (A-B) Two polyclonal cell populations targeted with HUWE1 sgRNAs (1 and 2) and one polyclonal cell population targeted with SCR sgRNA were derived for each genotype as described in Materials and methods. (A) Immunoblots of total HUWE1 from WCE used for quantification in B. The “a” and “b” superscripts next to the protein names indicate which of two duplicate membranes the corresponding strips were cut from. Dashed vertical lines indicate a rearrangement of samples within the same blot. (B) HUWE1 abundance (average HUWE1 intensity normalized to either Na^+^/K^+^ ATPase or GAPDH intensity from duplicate immunoblots shown in A) in WCE of cell populations targeted with HUWE1 sgRNAs, reported as percentage of HUWE1 abundance in WCE of cell populations targeted with SCR sgRNA control.

**S6 Fig.**
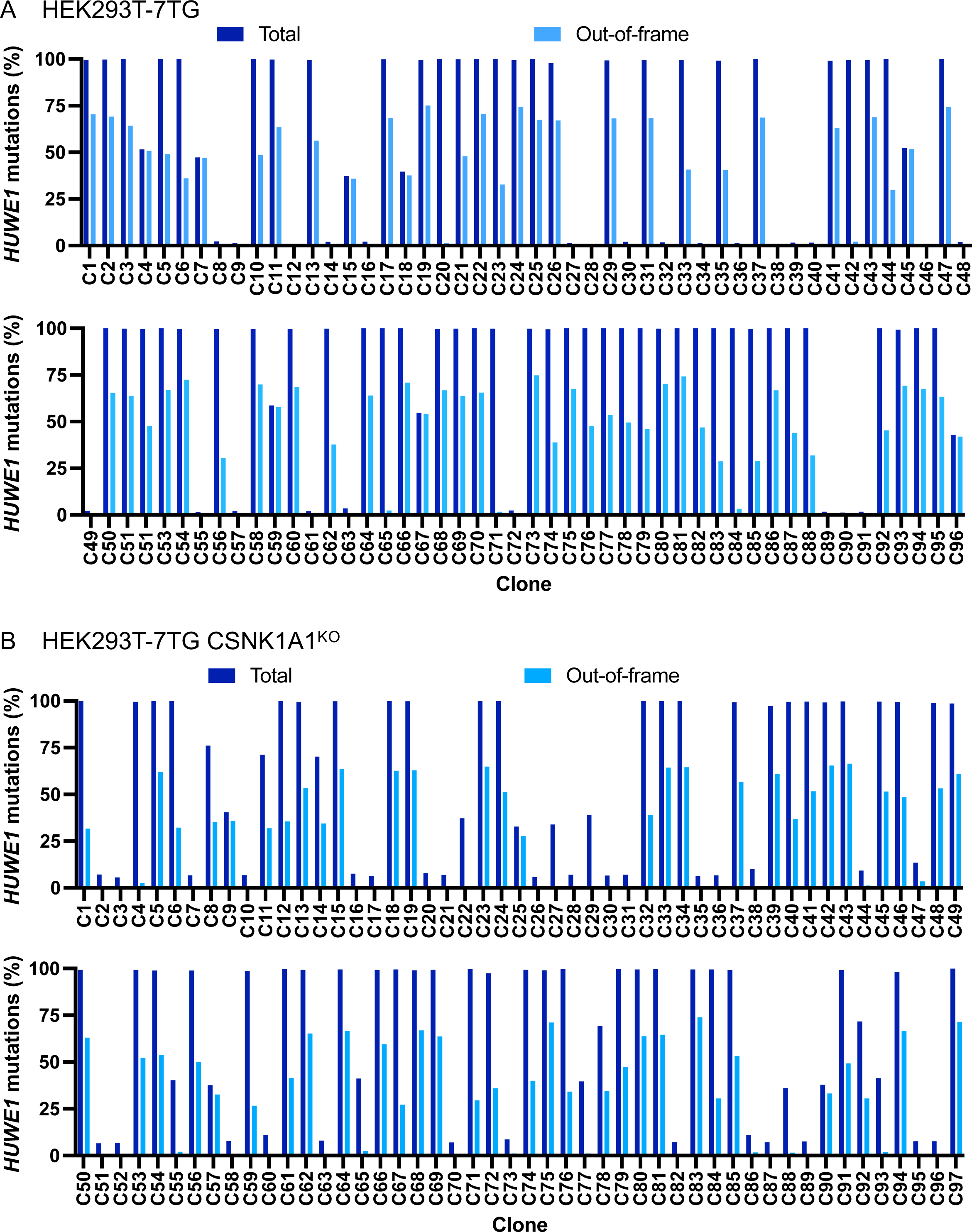
Quantification of CRISPR/Cas9-mediated *HUWE1* mutations in HEK293T-7TG and HEK293T-7TG CSNK1A1^KO^ cells. (A, B) Sequencing reads of the *HUWE1* locus targeted by CRISPR/Cas9 in individual clonal cell lines derived from HEK293T-7TG (A) or HEK293T-7TG CSNK1A1^KO^ (B) cells were quantified for mutations. The X-axis shows individual clones, and the Y-axis indicates the percentage of reads containing mutations. Bars in dark blue indicate the percentage of reads containing any kind of mutation (total mutations) at the targeted locus in each clone, and bars in light blue indicate the percentage of reads containing out-of-frame mutations at the same locus. In all 113 clones in which ∼100% of the reads contained mutations (indicating all *HUWE1* alleles had been successfully targeted), some of those mutations were always in frame, strongly suggesting that at least one WT *HUWE1* allele is required for cell viability in HEK293T cells.

## Supporting Information

**S1 File. CRISPR/Cas9-engineered clonal cell lines used in this study.**

Single-mutant clones in which a single gene was targeted using CRISPR/Cas9 and double- or triple-mutant clones in which multiple genes were targeted using CRISPR/Cas9 are described in two separate spreadsheets labeled accordingly. When more than one clone was generated using the same CRISPR guide, the ‘Clone Name’ column indicates the generic name used throughout the manuscript to describe the genotype, and the ‘Clone #’ column identifies an individual clone. The ‘HDR Donor’ column indicates the name of the ssODN donor template used to generate some of the clonal cell lines (see Materials and Methods). The ‘CRISPR guide’ column indicates the name of the guide used, which is the same as that of the oligos encoding sgRNAs (see Materials and methods, and S2 File). The ‘Genomic Sequence’ column shows 80 bases of genomic sequence (5’ relative to the gene is to the left) surrounding the target site. For each group of clones made using the same CRISPR guide (separated by gray spacers), the ‘Genomic Sequence’ column is headlined by the reference WT genomic sequence (obtained from RefSeq), with the guide sequence colored blue. The site of the double strand cut made by Cas9 is between the two underlined bases. Sequencing results for individual clones are indicated below the reference sequence. Some clones that remained WT at the targeted locus are indicated as such and were used as controls. For mutant clones, mutated bases are colored red (dashes represent deleted bases, three dots are used to indicate that a deletion continues beyond the 80 bases of sequence shown, and large insertions are indicated in brackets), and the nature of the mutation and the resulting genotype are described in the columns labeled accordingly. The figures in which each clone was used are also indicated. For double- and triple-mutant clones, the CRISPR guide used, the genomic sequence, the mutation and the genotype pertaining to each of the two or three targeted loci are designated ‘1’, ‘2’ and ‘3’ in the column headings, and are shown under green, orange and purple spacers, respectively.

**S2 File. Oligonucleotides and primers used in this study.**

Oligonucleotides and primers used for generation and characterization of clonal cell lines engineered using CRISPR/Cas9 nuclease (CRISPRn), base editing, and CRISPRi, as well as those used for qRT-PCR, are described in separate spreadsheets labeled accordingly. CRISPRn, base editing and CRISPRi: the names and sequences of pairs of oligonucleotides encoding sgRNAs, which were cloned into the respective vectors for each application as described in Materials and methods, are shown in columns A and B, respectively. Additionally, for CRISPRn and base editing the names and sequences of pairs of forward and reverse primers used to amplify corresponding genomic regions flanking sgRNA target sites are shown in columns C and D, respectively, and where applicable, the names and sequences of individual primers used to sequence the amplified target sites are shown in columns E and F, respectively. qRT-PCR: the names and sequences of pairs of forward and reverse primers used for qRT-PCR are shown in columns A and B, respectively.

